# Identification and characterization of new *P. aeruginosa* phages using a surface receptor platform

**DOI:** 10.64898/2026.11.06.764861

**Authors:** Veronica N. Tran, Tanisha S. Lahane, Ikram Qaderi, Hanjeong Harvey, Lori L. Burrows

## Abstract

Bacteriophages are of increasing interest for their potential use as alternatives to antibiotics for treating drug-resistant infections. Phage steering, specifically receptor-based steering, involves use of phages that bias the development of resistance in bacteria towards phenotypes such as reduced fitness. Phage therapy typically involves administering cocktails of phages with different targets to reduce the likelihood of resistance emerging. To ensure effective steering and cocktail design, there is a need to identify the receptors used by phages of interest, and to isolate phages that use diverse receptors. Here we describe a SURFAce Receptor (SURFAR) platform consisting of a set of defined *Pseudomonas aeruginosa* strains and mutants for rapid characterization of phage receptors. This platform can also be used to enrich for and isolate diverse phages from environmental samples, yielding novel phage genomes from wastewater. Using the SURFAR platform, two new phages were isolated and characterized: Dalewood, which targets both flagella and the conserved LPS outer core, and Bowman, a biofilm targeting phage capable of supressing growth of a subset of carbapenem-resistant clinical isolates. This tool helps to standardize the laborious but important task of phage receptor characterization, with the goal of expanding the known repertoire of *P. aeruginosa* phage receptors and phages.

**SIGNFICANCE:** Partially guided by increasing interest in phage therapy and technological advances, novel phages are regularly being discovered. Identification of phage receptors is an important aspect of initial characterization, but this step can be difficult due to lack of standardized strains and methodology. Most *P. aeruginosa* phages target the type IV pilus, LPS O-antigen, or flagella, thus there is a need to find phages with more diverse receptors. SURFAR was successfully used to identify the host ranges of phages with known receptors, data that can act as a reference during future characterization of new phages. Many SURFAR strains lack those canonical structures to bias for selection of phages that recognize new *P. aeruginosa* receptors. We showed that enrichment of wastewater filtrates using SURFAR strains yielded novel phage genomes, including new phages with therapeutic potential. Receptor-led characterization and isolation strategies with the SURFAR platform is a promising approach for finding new phages that can be combined into therapeutic cocktails of phages that target different receptors to mitigate resistance.

## INTRODUCTION

Antibiotic-resistant infections continue to rise each year, contributing to increased mortality and burden on health systems (1). As a result, there is a renewed interest in bacteriophages (phages) as treatments or adjuvants to current antimicrobial strategies. Prior to the age of antibiotics, phages (viruses that only infect bacteria) were used to treat bacterial infections. Antibiotics later replaced phages as preferred therapeutics because phages have a narrow host range and a high risk of biocontamination (2, 3). As a result, phages were broadly abandoned as therapeutics in Western countries (4), but remained a key scientific tool to study bacteria.

Phages are highly specific, often displaying strain-level specificity (5, 6). This selectivity is mediated, in part, by the first step of phage replication: adsorption to a cell-surface phage receptor (6). To avoid phage infection, bacteria can lose, downregulate, and/or modify the phage receptor (7–9). Phages may then fail to infect the cells or evolve to recognize the modified receptor, fueling an evolutionary arms race (10, 11). Since this inevitable back-and-forth can decrease therapeutic efficacy, phages targeting different receptors are typically administered as a cocktail to delay resistance (12, 13). While some phages in these cocktails have known receptors, phages with unknown or unreported receptors have also been administered to patients (9).

*Pseudomonas aeruginosa* is a Gram-negative, opportunistic pathogen listed on the World Health Organization’s high priority list and classified as an ESKAPE pathogen, emphasizing the need for novel approaches to combat treatment-refractory cases (14, 15). Most characterized *P. aeruginosa* phages recognize one of three common receptors: the O-specific antigen (OSA) O-antigen of the lipopolysaccharide (LPS), type IV pili (T4P), or flagella. The limited number of receptors and lack of detailed receptor reporting for many *P. aeruginosa-*targeting phages impede the rational receptor-based design of phage cocktails.

Modest receptor diversity is partly the result of isolation of phages using wild-type (WT) strains that express the most common receptors. Further, phages are often isolated from wastewater samples and enriched with WT strains in rich media, therefore the use of limited sources and media types may also contribute to reduced diversity.

The common receptors contribute to bacterial fitness and are therefore useful phage steering targets, but they are not conserved across strains. *P. aeruginosa* expresses at least 20 different OSA O-antigens, and O-antigen-targeting phages are often serotype specific (16). There are two different flagellin groups and five different pilin groups, with additional micro-diversity within groups (9). Pilin diversity is predicted to result from the selection of randomly arising pilin variants that mediate escape from phage infection and the group specificity of some T4P-targeting phages is consistent with this hypothesis (7, 17). Flagellins and pilins may be further masked by post-translational modifications and O-antigens may undergo serotype switching, both of which can prevent binding of phages that target the original variant (7, 9, 18–21). Thus, when using phages that recognize these receptors, the specific variant expressed by the target strain or isolate must first be identified to ensure compatibility with the phage. Using phages that recognize receptors that are more widely conserved among *P. aeruginosa* strains has the potential to reduce incompatibility.

With the goal of expanding receptor diversity of *P. aeruginosa* phages, and a focus on receptors that are likely to be conserved in multiple strains, we developed the SURFAce Receptor (SURFAR) platform. It consists of a curated, defined, and expandable set of WT, mutant, and recombinant strains for identification of phages that bind conserved, non-canonical receptors. These non-canonical receptors can be virulence factors, surface components that are clinically relevant for infection, and/or homologs of reported phage receptors in other species of bacteria. SURFAR can also be used to characterize the host range and probable receptor(s) of novel phages. This platform has the potential to produce useful host-receptor information that is applicable for receptor-led curation of phage cocktails and for training of large learning models. Using SURFAR, two new phages were identified and characterized. One of which, Bowman, is a biofilm matrix-targeting phage that can kill carbapenem-resistant clinical isolates, demonstrating its therapeutic potential.

## RESULTS

### Creation of the SURFAR platform

Strains included in SURFAR were selected based on their defined pilin groups, flagellin types, degrees of LPS modification, plus the loss or overexpression of potential receptors. Strains are summarized in **Table 1** and were generated as described in Methods. Notably, phages may fail to infect a WT strain despite the presence of a functional receptor, since replication may be halted by downstream anti-phage systems frequently found in clinical and environmental isolates (22). For this reason, many mutants in SURFAR are on the PAO1 strain background which is CRISPR-Cas deficient, meaning phage resistance is more likely attributable to receptor incompatibility (23). In addition to PAO1, four other common laboratory strains and their isogenic non-piliated mutants were included, representing the five pilin groups (PA1244-group I, PAO1-group II, PA14-group III, PA5196-group IV, and PA1457-group V) (17, 24). There are three strains with flagellum-related mutations: PAO1 *fliC* and PAK *fliC* lack the two types of flagellin (PAK: a-type; PAO1: b-type) (19). PAO1 *flgE* lacks the flagellum hook protein, a surface exposed component of the flagellar structure that is involved in infection and replication of *Salmonella enterica* targeting phage Chi (25).

**Table 1.** List of strains in the SURFAR platform.

| Receptor | Group | Strain |
|---|---|---|
| ***MUTANT STRAINS*** |  |  |
| **WT Reference** | Wildtype | PA1244 |
| **WT Reference** | Wildtype | PAO1 |
| **WT Reference** | Wildtype | PA14 |
| **WT Reference** | Wildtype | PA5196 |
| **WT Reference** | Wildtype | PA1457 |
| **Pilus** | Loss of potential receptors | PA1244 NP |
| **Pilus** | Loss of potential receptors | PAO1 NP |
| **Pilus** | Loss of potential receptors | PA14 NP |
| **Pilus** | Loss of potential receptors | PA5196 NP |
| **Pilus** | Loss of potential receptors | PA1457 NP |
| **Flagellum** | Loss of potential receptors | PAO1 *fliC* |
| **Flagellum** | Loss of potential receptors | PAK *fliC* |
| **Flagellum** | Loss of potential receptors | PAO1 *flgE* |
| **LPS** | Loss of potential receptors | PAO1 *wbpH* |
| **LPS** | Loss of potential receptors | PAO1 *wzm* |
| **LPS** | Loss of potential receptors | PAO1 *ssg* |
| **LPS** | Loss of potential receptors | PAO1 *galU* |
| **Canonical combo** | Loss of potential receptors | PAO1 *fliC/pilA* |
| **Canonical combo** | Loss of potential receptors | PAO1 *ssg/pilA* |
| **Canonical combo** | Loss of potential receptors | PAO1 *ssg/pilA/fliC* |
| **Canonical combo** | Loss of potential receptors | PAO1 *pilA/fliC* |
| **TBDT** | Loss of potential receptors | PAO1 *tonB1* |
| **TBDT** | Loss of potential receptors | PAO1 *tonB2* |
| **TBDT** | Loss of potential receptors | PAO1 *fpvA* |
| **TBDT** | Loss of potential receptors | PAO1 *fpvB* |
| **TBDT** | Loss of potential receptors | PAO1 *fptA* |
| **Outer-membrane proteins** | Loss of potential receptors | PAO1 *oprF* |
| **Outer-membrane proteins** | Loss of potential receptors | PAO1 *oprJDM* |
| **Regulator** | Loss of potential receptors | PAO1 *mexT* |
| **Biofilm negative** | Loss of potential receptors | PAO1 *pelA* |
| **Biofilm negative** | Loss of potential receptors | PAO1 *psl* |
| ***OVER-EXPRESSION STRAINS*** |  |  |
| **TBDT overexpression** | Overexpression of potential receptors | PAO1 +pB-*fpvA* |
| **TBDT overexpression** | Overexpression of potential receptors | PAO1 +pB-*fpvB* |
| **TBDT overexpression** | Overexpression of potential receptors | PAO1 +pB-*fptA* |
| **TBDT overexpression** | Overexpression of potential receptors | PAO1 +pB-*foxA* |
| **TBDT overexpression** | Overexpression of potential receptors | PAO1 +pB-*piuA* |
| **TBDT overexpression** | Overexpression of potential receptors | PAO1 +pB-*fiuA* |
| **Lacking canonical receptors TBDT overexpression** | Overexpression of potential receptors | PAO1 *pilA/wbpH/wzm* +pB-*fpvA* |
| **Lacking canonical receptors TBDT overexpression** | Overexpression of potential receptors | PAO1 *pilA/wbpH/wzm* +pB-*fpvB* |
| **Lacking canonical receptors TBDT overexpression** | Overexpression of potential receptors | PAO1 *pilA/wbpH/wzm* +pB-*fptA* |
| **Lacking canonical receptors TBDT overexpression** | Overexpression of potential receptors | PAO1 *pilA/wbpH/wzm* +pB-*foxA* |
| **Lacking canonical receptors TBDT overexpression** | Overexpression of potential receptors | PAO1 *pilA/wbpH/wzm* +pB-*piuA* |
| **Lacking canonical receptors TBDT overexpression** | Overexpression of potential receptors | PAO1 *pilA/wbpH/wzm* +pB-*fiuA* |
| **Biofilm hyper-producer** | Overexpression of potential receptors | PAO1 +pB-*sadC* |
| **Biofilm hyper-producer** | Overexpression of potential receptors | PAO1 *pilA/wbpH/wzm* +pB-*sadC* |
| **Biofilm hyper-producer** | Overexpression of potential receptors | PAO1 *fliC/pilA* +pB-*sadC* |

pB= pBADGr, WT= wildtype, LPS= lipopolysaccharide; see **Supplementary Table S2** for strain sources and reference

Four LPS-related mutants on the PAO1 background were included. LPS consists of membrane-embedded lipid A, inner- and outer-core sugars, and long-chain O-antigen attached to a subset of those cores (16). *P. aeruginosa* simultaneously expresses two forms of LPS O-antigens: the common polysaccharide antigen (CPA) is a homopolymer of D-rhamnose, and the OSA is a heteropolymer whose sugar compositions, glycosidic linkages, and lengths depend on the specific serotype (16). Most known LPS-targeting phages recognize OSA O-antigens, therefore we have included PAO1 *wbpH* which causes loss of OSA (26). PAO1 *wzm* lacks the highly conserved CPA O-antigen (27). The products of *galU* and *ssg* are involved in outer core subunit metabolism and synthesis, thus the conserved inner core is exposed in strains with these mutations (28).

Mutants devoid of select outer membrane proteins were included, based on at least one of the following criteria: they are *P. aeruginosa* orthologs of porins and surface proteins identified as phage receptors in other bacterial species with predicted surface exposure, their potential roles in fitness or virulence that can be leveraged in steering, and/or they are highly conserved amongst *P. aeruginosa* strains (29–31). For example, OprF is the most abundant porin in *P. aeruginosa* and facilitates the general transport of large solutes (32). It also contributes to maintenance of cell architecture, as *oprF* mutants display cell morphology defects. OprF is also suggested to have a role in biofilm matrix formation (31, 33, 34). OprM is the outer membrane transporter component of the most abundant efflux systems: MexAB-OprM and MexXY-OprM, which are responsible for extrusion of antibiotics (35, 36). *P. aeruginosa* phage OMKO1 was suggested to recognize OprM (30). OprJ is the outer-membrane component of MexCD-OprJ system and is also responsible for antibiotic efflux, contributing to fluoroquinolone resistance (37, 38). MexT is a transcriptional regulator of various systems including positive regulation of MexEF-OprN, negative regulation of OprD, and MexS, which in turn negatively regulates MexT (39–41). Thus, *mexT* mutants may select for phages that depend on MexEF-OprN but not OprD or other MexS dependent factors for infection. These systems are involved in efflux of different antibiotics to contribute to development of resistance, and the mutants lack either a receptor or a regulator of receptor expression. Using phages that target these potential receptors may steer bacteria towards loss of the receptor, disabling these AMR strategies.

TonB-dependent transporters (TBDTs) are gated outer membrane channels whose opening is powered by TonB1 and/or TonB2 to allow for uptake of iron chelating siderophores (42, 43). These components are upregulated in low iron environments, including in the human host (43). When iron bound siderophores interact with one of the transporters, a conformational change leads to an interaction between the transporter and TonB. This triggers an inner membrane complex of TonB-ExbB-ExbD to leverage the proton motive force to power active transport of the chelated iron into the cell (43–46). TonB1 is the primary energizer, while TonB2 is activated in specific conditions; the two can partially cross complement. Loss of both proteins prevents energization of the TBDTs, a requirement for phage replication in other systems (47). Different TBDTs are specific for uptake of different chelators including xenosideophores (siderophores encoded by other species) and various siderophores secreted by the cell itself (46). Certain antibiotics may also use TBDTs to enter the cell, and orthologs of TBDT are phage receptors in other species of bacteria (46, 48). Phages that infect in a TonB-dependent manner may steer bacteria to loss of the transporters and an inability to uptake iron, impacting fitness in low iron environments. Some mutants lack more than one receptor, such as PAO1 *fliC/pilA* which is devoid of both flagella and pili, or PAO1 *tonB1* and PAO1 *tonB2* which powers active transport of multiple TBDTs, therefore these mutations inactivate all those transporters (42, 49, 50).

Mutants lacking known and potential receptors were created either through transposon insertion in the target gene (51), whole gene deletion through splicing by overlap extension PCR (52), or FRT insertion (53), followed by confirmation by sequencing. Some strains reached a higher optical density (OD) and stationary phase more quickly than others, suggesting variations in growth patterns between the strains, but all were capable of phage propagation. Of note, strains overexpressing *sadC* and those with LPS core mutations (*ssg* and *galU*) required longer growth times to reach the same OD as their counterparts. *P. aeruginosa* clinical isolates can display growth defects compared to lab strains, so the platform is representative of the overall fitness levels of *in vivo* bacterial populations.

The platform also contains strains overexpressing potential receptors to increase the opportunity of phage interaction. These potential receptors are not highly expressed in the rich media used in our phage enrichment experiments, therefore *in trans* overexpression bypasses this hurdle. To create overexpression strains, genes encoding select TBDTs were inserted into an arabinose inducible plasmid (54) to allow for *in trans* titration of receptor expression (46). SadC is a diguanylate cyclase that promotes synthesis of cyclic di-GMP, a secondary messenger that increases biofilm formation in *P. aeruginosa* through increased expression of biofilm matrix components such as polysaccharides (55, 56). To create biofilm-hyperproducing strains, *sadC* was overexpressed *in trans* in strains WT PAO1, PAO1 *pilA/wbpH/wzm (*PAO1 AHM) which lacks T4P and both OSA and CPA O-antigens, and PAO1 *fliC/pilA* which lacks the flagellum and T4P. All strains were organized into a 96 deep well plate format.

### Validation of the SURFAR platform for phage host and receptor characterization

With the growing number of newly identified phages (57), there is a need to standardize the properties assessed during characterization. While genome sequence, morphology, and host range are consistently documented, receptor usage is inconsistently reported despite its central role in phage-host interactions (9). This gap stems partly from a lack of consensus strains or methods to use for receptor characterization. To address this limitation, we propose that SURFAR can be used to help identify the receptor(s) of a given phage.

To validate this application of SURFAR, we assessed the infectivity patterns of six previously characterized lytic *P. aeruginosa* phages with known receptors (58, 59) (**Figure 1**). These phages can provide reference data when using SURFAR to characterize novel or previously uncharacterized phages, as together they span host and/or receptor profiles representing commonly characterized receptors that are also of clinical relevance.

**Figure 1.**
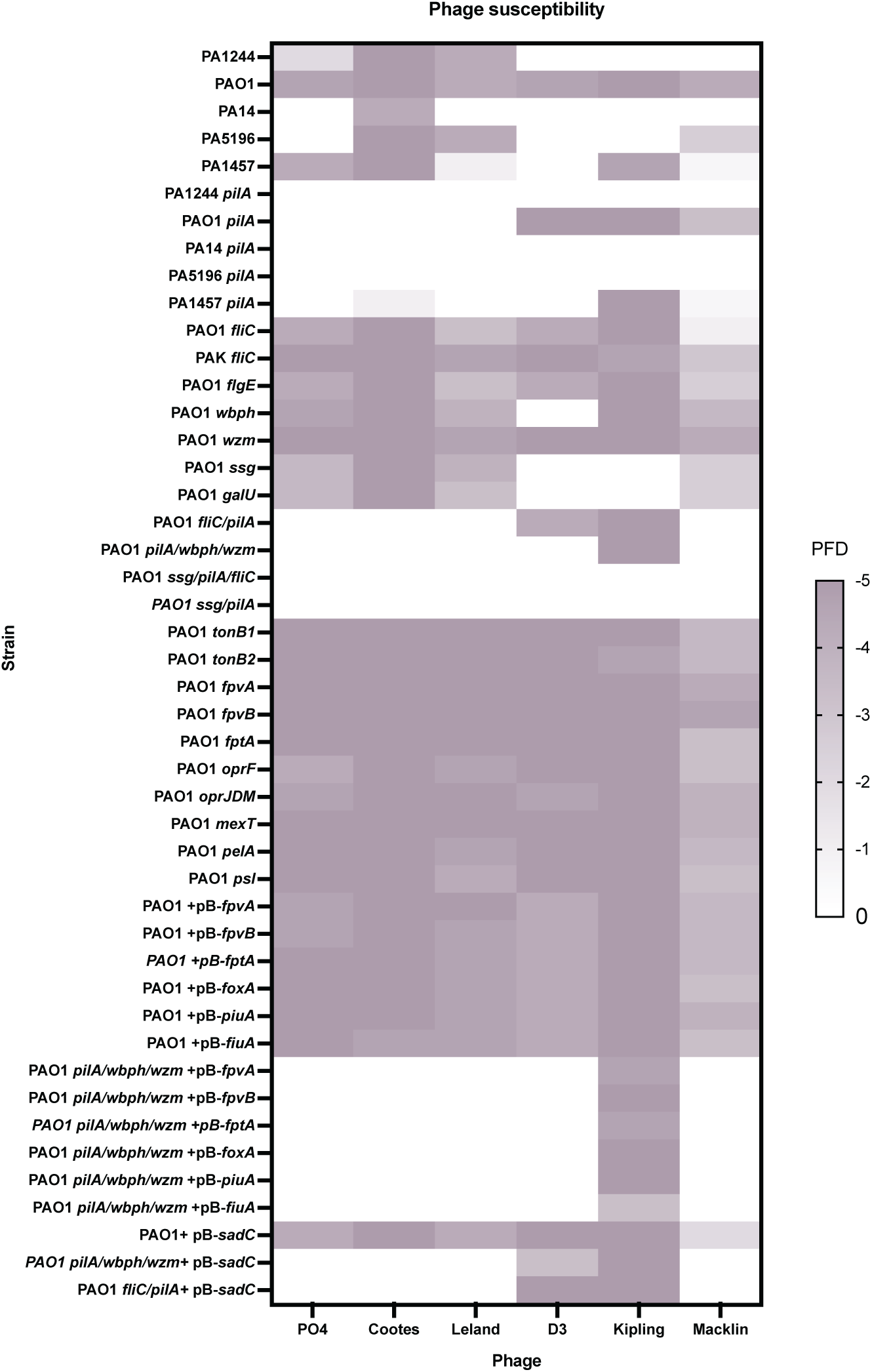
Plaquing patterns of characterized phages are consistent with previously reported host ranges and receptor profiles. Phages PO4, Cootes, Leland, D3, Kipling, and Macklin were plaqued on the SURFAR platform. Strains of platform are indicated on the left. Darker colour indicates higher plaque forming dilutions (PFD) with white indicating no plaques/infection. pB=pBADGr, PFD= dilution where single plaques are present. Data is representative of three independent experiments.

PO4 is a well-characterized phage that recognizes the T4P (8, 58). Cootes infects in a pilus-dependent manner and recognizes all five pilin groups, while phage Leland recognizes four of the five pilin groups (60). D3 is another well-characterized phage that recognizes a single O-antigen serotype (18). While the LPS O-antigen is highly variable, the LPS outer core is conserved (16, 28). Phage Kipling uses the outer core as its receptor (60). Some phages require multiple receptors for infection: initial attachment to a primary receptor and irreversible attachment to a secondary receptor to initiate replication (61, 62). Phage Macklin recognizes both the pilus and flagella, and serves as a reference for other multi-receptor phages.

PO4 infected all PAO1-based strains expressing group II pilins, plus strains PA1244 and PA1457 which express groups I and V pilins, respectively. Its inability to plaque on the other WT strains could reflect pilin incompatibility or downstream phage defence mechanisms. Cootes infected strains expressing each of the five pilin groups. Pilus-deficient strains were resistant with the exception of PA1457 *pilA* where plaques were visible only at the highest dilution and lacked the clear plaque morphology seen in WT strains. Leland, the other broad pilus group infecting phage, infected all strains except for PA14 and pilus-deficient strains. Its inability to infect PA14 may be due to pilin group incompatibility or downstream anti-phage mechanisms (5, 24, 63, 64).

D3 infected only PAO1 and was unable to form plaques on strains lacking OSA, consistent with D3’s specificity for serotype O5 LPS (18, 59). Kipling infected only PAO1 strains and PA1457. It did not infect PAO1 strains carrying *ssg* or *galU* mutations which disrupt the LPS outer core, but maintained infectivity against strains with O-antigen mutations as the LPS outer core is intact. Macklin infected WT strains of PAO1, PA1457, and PA5196, plus the non-piliated mutants of PAO1 and PA1457, but not the non-piliated mutant of PA5196. In PAO1, both T4P and flagella were required for plaquing at the lowest dilution and flagellum-deficient strains showed reduced plaquing efficiency relative to PAO1 *pilA.* Macklin uses both receptors, with flagella likely serving as a secondary receptor. Both PAO1 and PA5196 express b-type flagellin (65), while the flagellin type of PA1457 is unknown. Susceptibility across these strains may be mediated by flagellin type compatibility and absence of downstream anti-phage strategies that would otherwise prevent infection and replication.

All the phages infected strains with mutations in TonB-related genes, porins, and *mexT*, suggesting that these components are not required for replication. These results show that SURFAR can identify phage receptors down to the level of strain-specific differences, as seen through PO4 pilin-groups and D3 serotype specificity.

### SURFAR selects for genetically diverse phages

In addition to characterization of purified phages, we propose that SURFAR can be used for the enrichment and isolation of new phages. SURFAR strains can be broadly categorized into five groups: WT strains of *P. aeruginosa* expressing different pilin types, flagellin types, and O antigen serotypes (purple in Figures 2-4); strains lacking a single characterized receptor (blue); strains devoid of multiple known receptors; (green) strains lacking predicted receptors (orange/yellow); and strains overexpressing predicted receptors (red/pink). We hypothesized that overexpressing predicted receptors in a WT background would increase the likelihood of phage interaction. Additionally, because these predicted receptors may serve as secondary receptors, expressing them in strains with the canonical primary receptors ensures that phages have access to both. Alternatively, canonical receptors may mask alternate receptors, so we also overexpressed potential receptors in PAO1 strains devoid of pili and both types of LPS O-antigens (*pilA/wbpH/wzm*).

**Figure 2.**
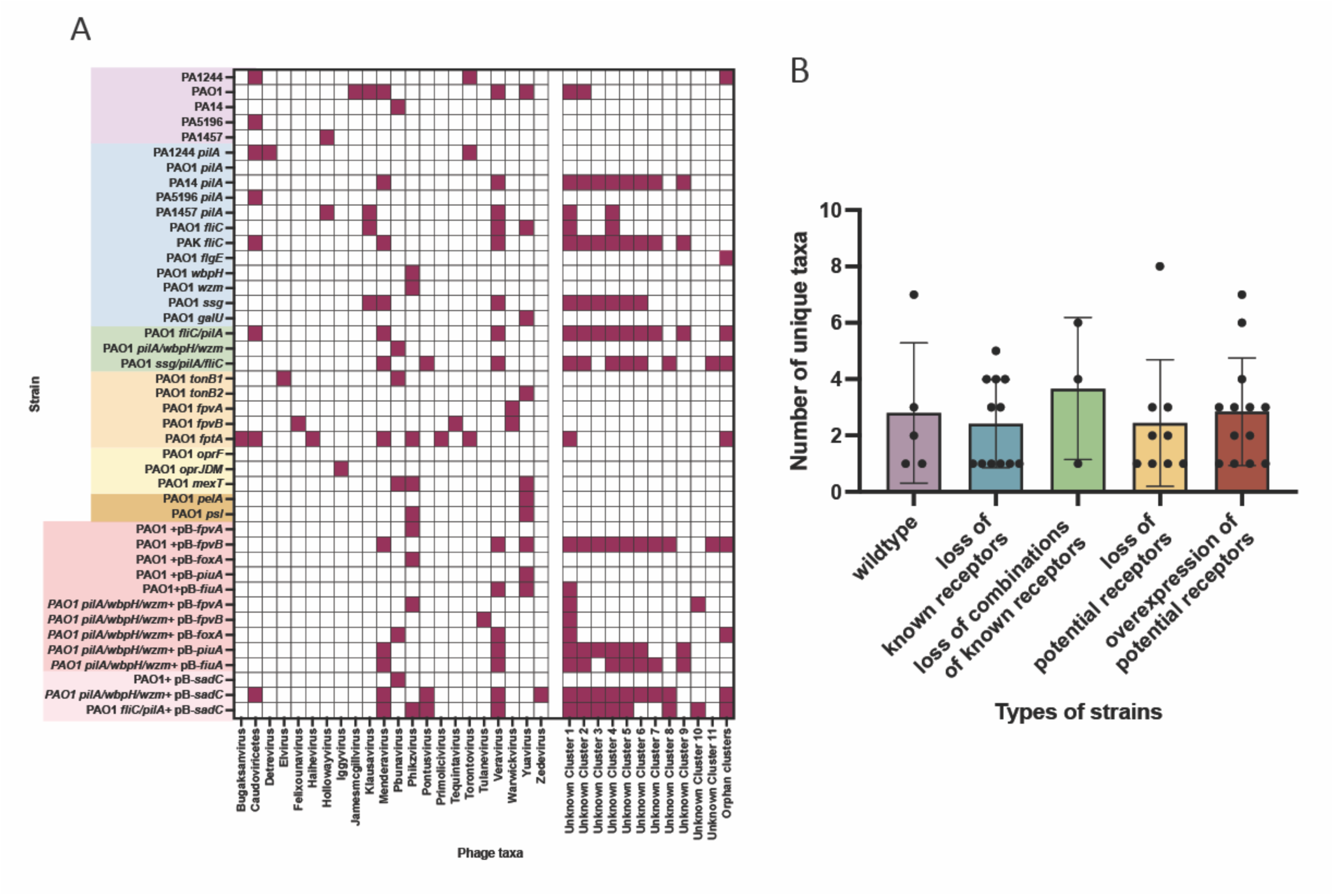
SURFAR strains enrich for diverse phage taxa in wastewater samples. **A)** Taxa of phages (bottom) in lysates are indicated according to the SURFAR strain (left) used to enrich wastewater sample filtrates. Purple indicates presence of phage contigs with homology to indicated taxa. Nine groups of related known contigs are indicated, contigs that had no homology to NCBI or INPHARED database taxa were clustered together based on sequence homology, with the final column containing orphan contig. SURFAR strains are colour coded according to the groups in **B),** which notes the average number of phages contigs from unique taxa in lysates enriched by each strain within the different enrichment strain groups. Value indicates mean, error bars indicate standard deviation. Data are from one round of enrichment.

Wastewater is a rich source of new phages (66, 67). Typically, influent samples from various treatment sites and collection dates is filtered to remove environmental debris and bacteria, then the filtered wastewater supplemented with LB media and enriched with a single *P. aeruginosa* strain to propagate any phages present, as described in the Methods. To increase the number and diversity of phages recovered in this study, filtered samples from four distinct wastewater collection sites and timepoints were combined in equal volumes prior to bacterial enrichment. Following incubation, phage-containing supernatants were collected, and in collaboration with Canada’s National Microbiology Lab’s PhageSTAR program, the lysates were sequenced to characterize phage contents.

A total of 367 contigs representing potential phage genomes were identified in 43 lysates. One lysate contained only bacterial DNA and was therefore excluded from further analysis. Of the putative phage contigs, 71% (259/367) had no matches in the NCBI BLAST or INPHARED databases and were classified as “unknown”. This suggested that many of the sequences are novel, as defined by their lack of homology to phage genomes in the databases. Only a subset of these contigs may represent *P. aeruginosa-*targeting phages, since phages that non-specifically bind *P. aeruginosa* enrichment strains, phage satellites, and free phage DNA fragments would all be captured in sequencing of lysates. The remaining 108 (29%) sequences had matches in the NCBI Viruses database and were classified into their respective taxa.

To evaluate whether any of the 259 unknown contigs were related, we compared their MASH scores and grouped those contigs with similar scores (MASH distance ≤ 0.05) as related or possible duplicate sequences (68). Of the unknown contigs, 97 clustered with at least one other contig and the remaining 162 are hereafter referred to as “orphan unknowns”. There was no clear association between the strain used for enrichment and the taxa or unknown clusters of the resulting contigs (**Figure 2A**). Some taxa were enriched by multiple strains, whereas others were enriched by only a few strains, and each group of strains selected for a similar number of taxa (**Figure 2B**). This suggests that increasing the diversity of isolated phages may depend less on the specific mutant or strain group used, compared to the number of enrichment and isolation rounds.

Since a large proportion of phage sequences lacked informative BLAST hits, taxonomic classification alone did not capture the total diversity of the samples, as 44% (162/367) of the contigs could not be clustered. Instead, we considered that evaluating phage sequence/contig diversity at the sequence level may provide a more comprehensive snapshot. Based on MASH distances, the 367 phage sequences could be organized into 22 distinct clusters with limited sequence similarity between clusters (**Figure 3A**) and many orphan clusters. This sequence classification is consistent with the taxonomic organization (**Figure 2A**).

**Figure 3.**
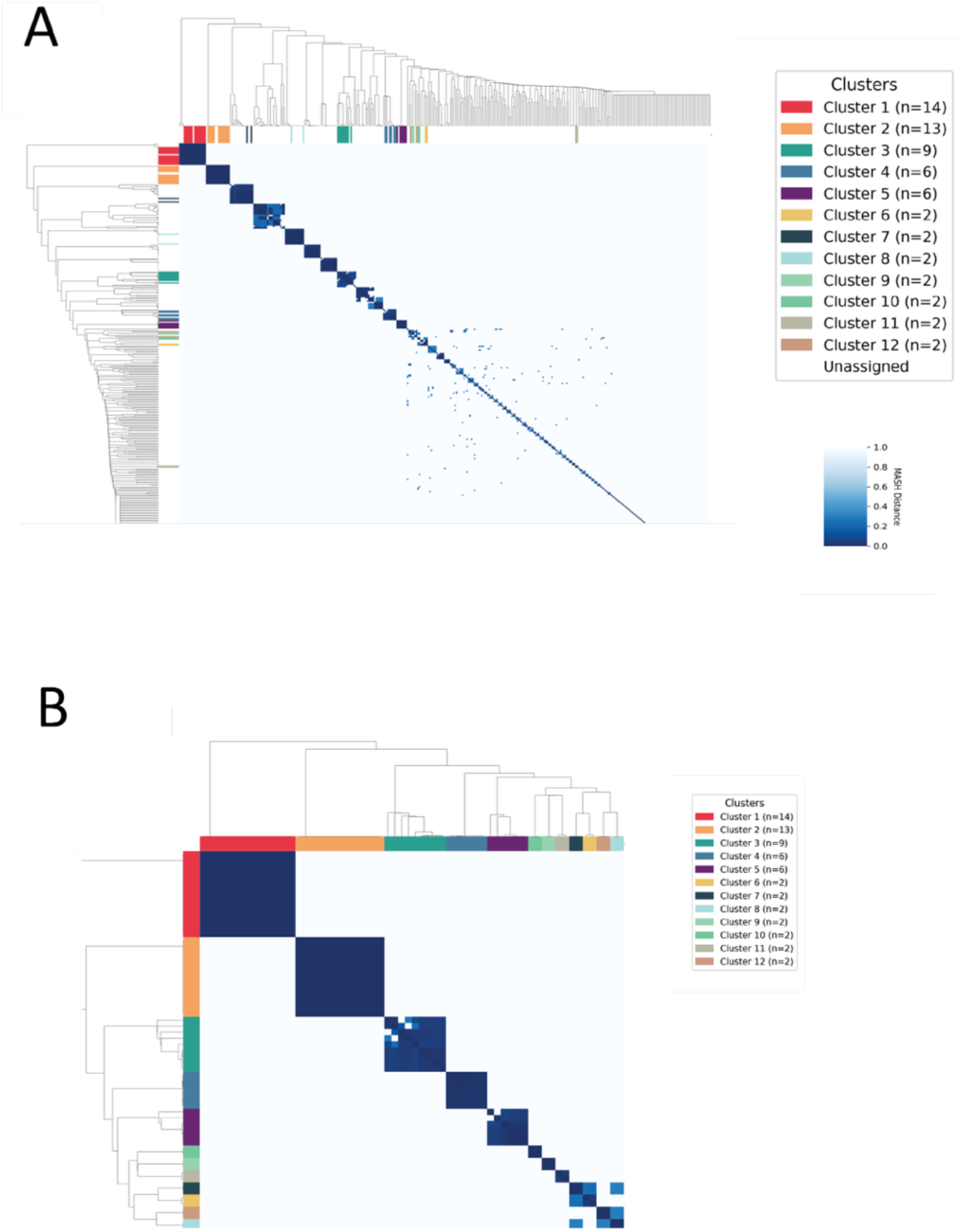
Genomic distance analysis of 367 phage contigs. **A)** Hierarchically clustered pairwise MASH distance matrix for 367 phage contigs assembled from SURFAR enrichment of wastewater samples. **B)** Heatmap of the 72 contigs assigned to high-confidence clusters (coloured clusters). Clustering threshold consisted of MASH distance <0.05, >20x sequencing coverage, and contig length > 15,000 bp. Dark blue indicates identical genomes while white indicates no detectable similarity. Clustering was performed and visualized using seaborn clustermap (v0.13.2).

**Figure 4.**
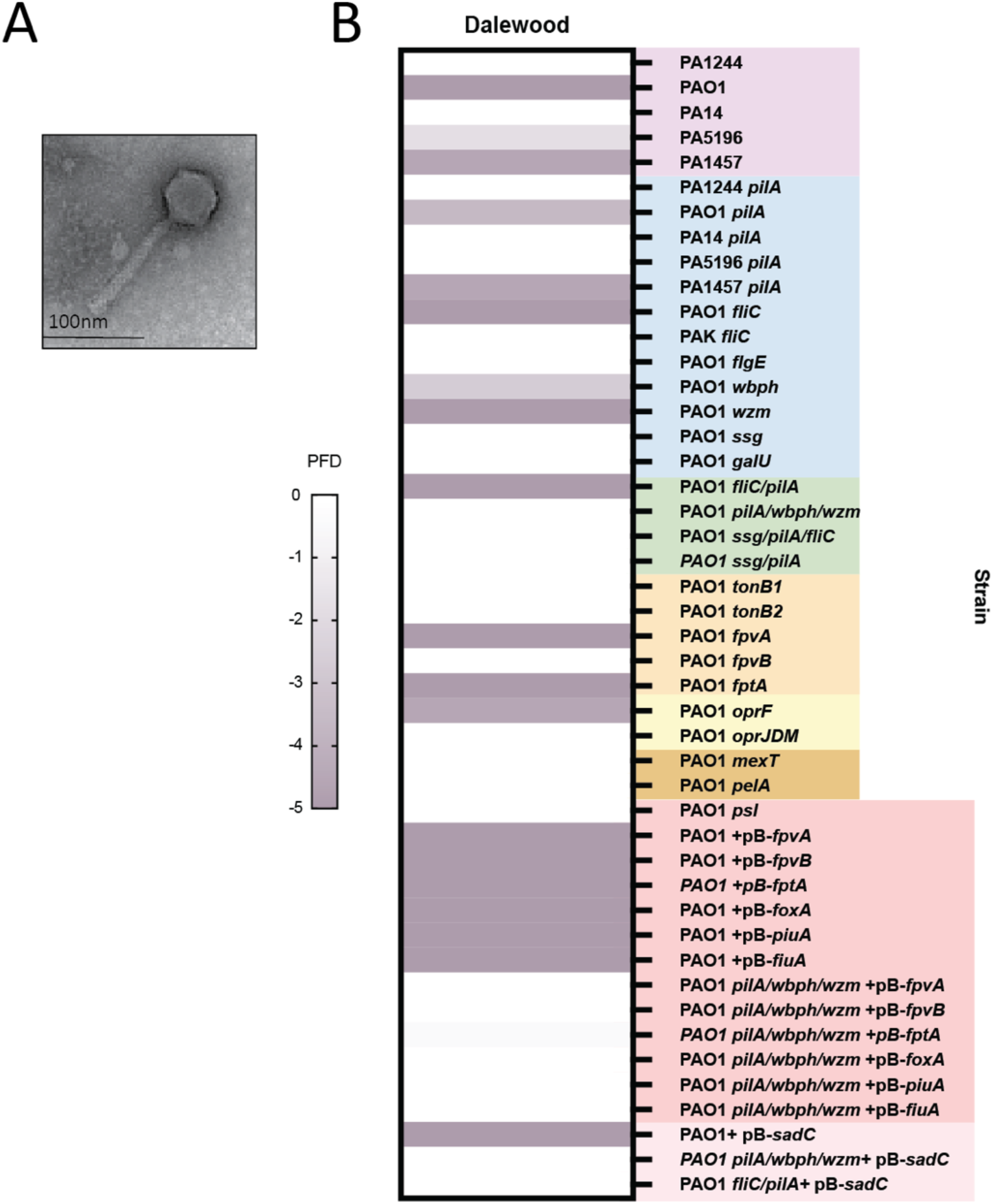
Characterization of phage Dalewood. **A)** Transmission electron microscopy image of Dalewood. **B)** Susceptibility of Dalewood to strains of the SURFAR platform (right). SURFAR strains are colour coded based on groups (purple=wildtype, blue= loss of known receptors, green = loss of combinations of known receptors, yellow/orange= loss of potential receptors, red/pink= overexpression of potential receptors). Darker colour indicates higher plaque forming dilution (PFD), with white indicating no plaques/infection. Data is representative of three independent experiments. pB=pBADGr, PFD= dilution where single plaques are present.

When a more stringent re-clustering criteria was applied, consisting of a minimum coverage cutoff of 20x and contig length of >15,000 bp, 295 contigs failed to meet this threshold, suggesting they may represent incomplete genomes. They were not included in secondary analysis. The remaining 62 genomes were further stratified into 12 clusters and there were no orphan clusters **(Figure 3B).** These data suggest that within the observed genomic diversity, some sequences may be duplicates or near-identical contigs of each other, as sequences with a MASH distance ≤ 0.05 were grouped together. This organization is in line with homology- and taxa-based classification of the contigs (**Figure 2A**).

We next examined whether there was a correlation between the sequence clusters (**Figure 3B**) and the strains used for enrichment. Due to the small sample size, statistical significance could not be calculated, therefore patterns described below are observational. There was no clear pattern among the strains used to enrich the first five clusters (**Supplementary Table S1**). Clusters 1 and 2, the largest clusters, were enriched using multiple *P. aeruginosa* strains and various PAO1 mutants. Both contigs are predicted to represent novel phages. Host prediction using vHULK and iPHoP were unsuccessful (69, 70) and these probable phages could not be plaque purified or isolated. Therefore, it remains unclear whether these contigs represent viable phages and what their hosts may be. Sequences in cluster 3 are all related to *Yuaviruses*; cluster 4 sequences all exhibit homology to *Pbunaviruses*, which typically use LPS as a receptor (71–73); and cluster 5 contigs are similar to the jumbo phage *Phikzvirus*. The remaining clusters contained fewer genomes and sequences within each individual cluster were enriched by similar strains (**Supplementary Table S1**).

To determine if the 259 unknown contigs represented phage genomes (incomplete or complete) and whether they might be from *P. aeruginosa* targeting phages, all contigs were annotated and the head protein and terminase large subunit (TLS) sequences were queried, if present. The head protein and TLS are among the most conserved proteins in phage genomes (74, 75). Both analyses returned numerous *Pseudomonas* phages and phage sequences from uncultured marine metagenomic samples (**Supplementary File S1**). Similar to wastewater, the marine environment is abundant in *Pseudomonads*, therefore these marine metagenomic samples likely contain *Pseudomonad-*targeting phages (76–78). In both analyses, the percent identity values were low and e-values were high, suggesting these sequences are distant homologs to well-conserved viral proteins, supporting the idea that most of the phages we enriched are truly novel (**Supplementary File S1**).

### Isolation of phage Dalewood

We next evaluated if SURFAR could be used to isolate and purify novel phages with clinical potential. Using the SURFAR platform, two new phages were isolated and characterized. One of these phages, Dalewood, belongs to the *Pbunavirus* genus (NCBI Accession: PZ594774). Transmission electron microscopy (TEM) imaging revealed an icosahedral head and tail morphology, similar to that reported for other phages in this genus (**Figure 4A**) (73, 79). Dalewood was isolated by enriching filtered wastewater samples with strain PAO1 Δ*wbpL* Δ*pilA,* which lacks the T4P and both O-antigens, followed by three rounds of plaque purification.

To characterize the host range and receptor(s) of Dalewood, SURFAR strains were challenged with the phage (**Figure 4B**). Dalewood could not infect strains carrying a mutation in *ssg*, which results in a truncated LPS core, suggesting that it likely uses LPS outer core its receptor. Dalewood could infect other strains of *P. aeruginosa,* including PA5196 and PA1457 which expresses different O antigen serotypes than PAO1. Interestingly, Dalewood did not infect a PA5196 pilin-deficient strain nor several strains in the “loss of potential receptors” category (PAO1 *tonB1, tonB2, fpvA, fptA, mexT, pelA, psl)*, suggesting these strains may carry an unidentified LPS core mutation.

### Isolation of biofilm-targeting phage Bowman

A mixture of three phages was isolated from wastewater enriched with *P. aeruginosa* PA1244. Among the three, the phage with the highest coverage belonged to the genus *Bruynoghevirus*. Following three rounds of plaque purification, this phage was sequenced, determined to be a new species, and named Bowman (NCBI Accession: PZ584235). TEM imaging of Bowman shows podovirus morphology consistent with other members of the genus (80, 81) (**Figure 5A**).

**Figure 5.**
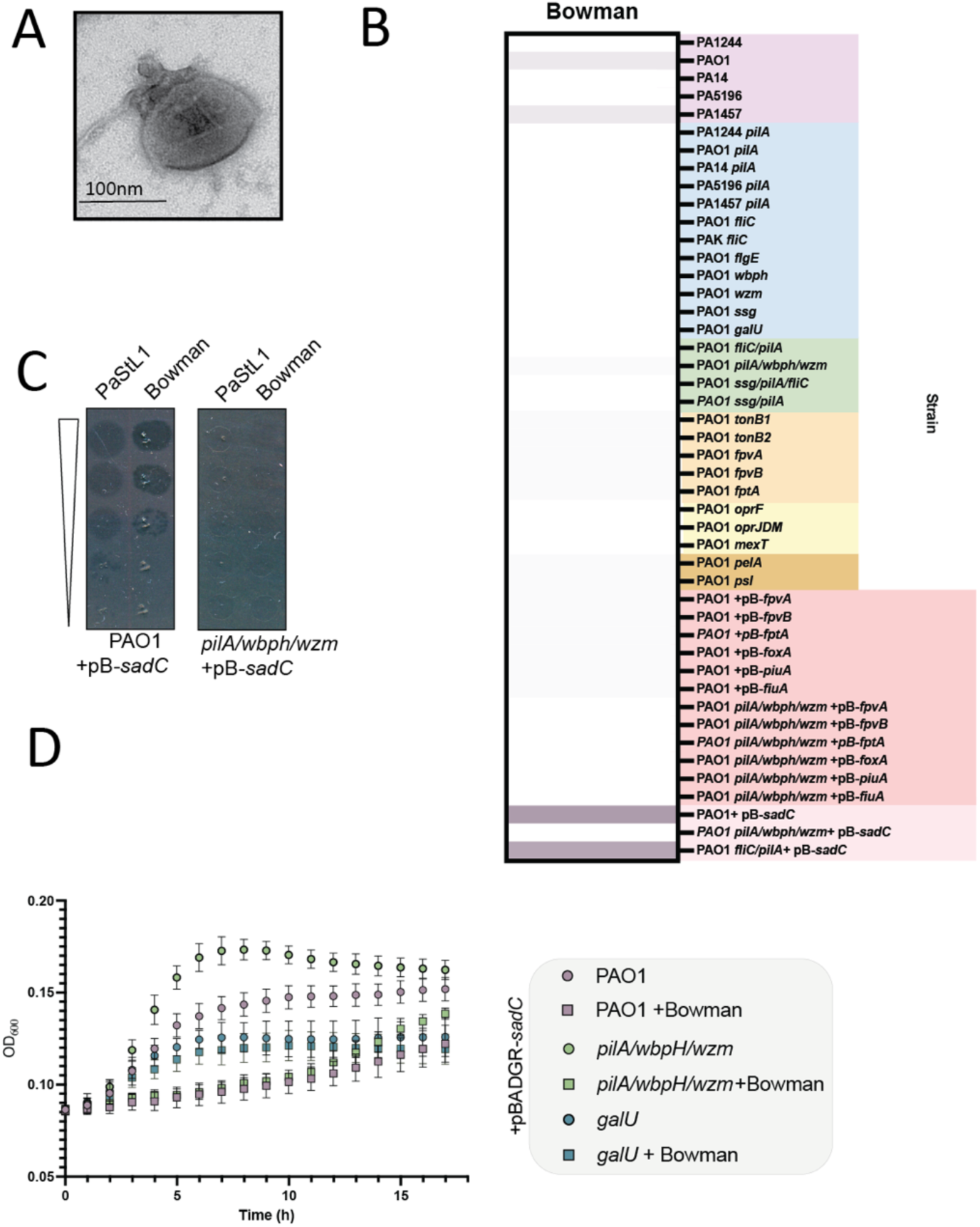
Characterization of phage Bowman. **A)** Transmission electron microscopy image of Bowman. **B)** Susceptibility of Bowman to strains of the SURFAR platform (right). SURFAR strains are colour coded based on groups (purple=wildtype, blue= loss of known receptors, green = loss of combinations of known receptors, yellow/orange= loss of potential receptors, red/pink= overexpression of potential receptors). Darker colour indicates higher plaque forming dilution (PFD). **C)** Bowman cannot form plaque on PAO1 *pilA/wbpH/wzm* but the strain is susceptible to Bowman in a **D)** liquid killing assay. Grey border indicates strains overexpressing *sadC.* Data is representative of three independent experiments. pB=pBADGr, PFD= dilution where single plaques are present.

*P. aeruginosa* typically grows as a biofilm *in vivo,* conferring protection against environmental stressors such as antibiotics and phages (82). The biofilm matrix consists of eDNA, proteins, lipids, and exopolysaccharides, including Psl (83). Psl is important for biofilm surface attachment, structural integrity, and immune evasion, and is the major expolysaccharide component in PAO1 (84). Novel *P. aeruginosa* phages that require Psl for infection were recently isolated and also belong to the *Bruynoghevirus* genus (80, 81). While various phages have been described to encode depolymerases that can degrade biofilm exopolysaccharides, only this genus of *P. aeruginosa* phages infects in a strictly biofilm-dependent manner (85–88).

SURFAR platform strains were challenged with Bowman to determine whether it is a biofilm exopolysaccharide-dependent phage, like other members of the genus, and if it requires additional receptors (**Figure 5B**. Bowman either did not plaque or formed plaques only at the highest titre on strains that were not hyper-biofilm producers. In contrast, it formed plaques beyond the first dilution on PAO1 + pB-*sadC* and PAO1 *fliC/pilA* + pB-*sadC* but not PAO1 AHM + pB-*sadC* (**Figure 5C**). Bowman could kill PAO1 AHM + pB-*sadC* in a liquid phage susceptibility assay (**Figure 5D**). This discrepancy between liquid killing and solid plaque assays has been observed in previous work from our lab and others (8). Bowman does not infect PAO1 *psl* or *pelA* mutants. These data suggest that Bowman, like other members of the genus (such as PaStL1 (81)), requires biofilms for infection.

We raised Bowman phage-resistant mutants (PRMs) on PAO1+ pB-*sadC.* Some colonies had a red phenotype associated with a genomic deletion encompassing *galU,* which encodes for UTP-glucose-1-phosphate uridylyltransferase, an enzyme involved in synthesis of the glucose subunits of the LPS core (28, 89, 90). Two other non-red PRMs were sequenced, also revealing mutations in LPS-related genes. One had a point mutation in *ssg* and the other had a 101 base deletion spanning three LPS-core biosynthesis genes: *dnpA,* PA4003, and *wapH.* The roles of ssg, *dnpA,* and PA4003 in LPS core biosynthesis are not well defined, but both *ssg* and *wapH* are predicted to contribute to the core architecture through the addition of glucose subunits (28). Therefore, we next evaluated the role of LPS in Bowman infection. Strains PAO1 Δ*galU* + pB- *sadC*, PAO1 Δ*ssg* + pB-*sadC*, and PAO1 *Δssg/ΔpilA/ΔfliC*+ pB-*sadC* strains were all resistant to Bowman (**Figure 6A**).

**Figure 6.**
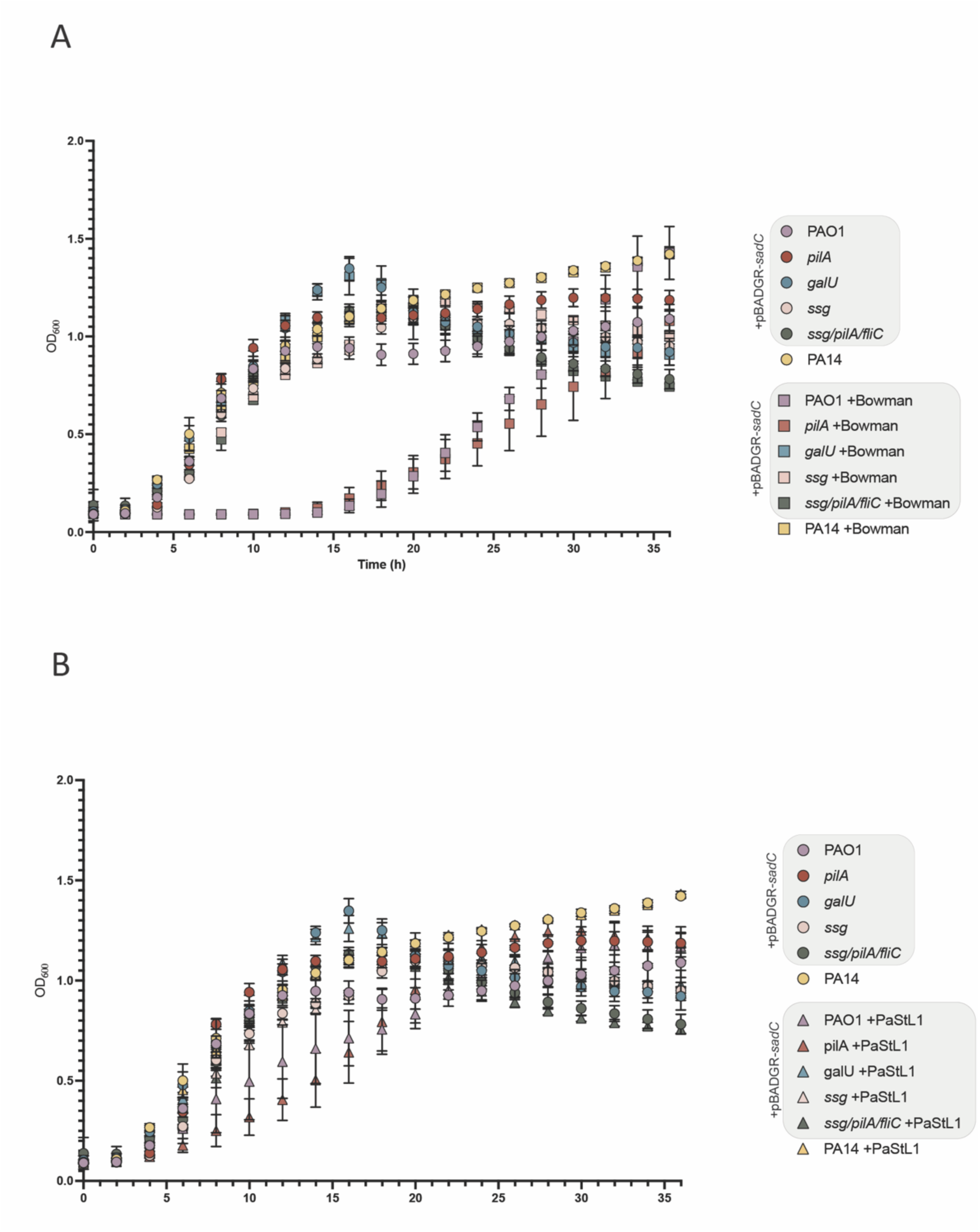
Bowman kills certain biofilm producing strains. **A)** Bowman can only infect PAO1 + pB-*sadC* and PAO1 *pilA* + pB-*sadC*, similar to the **B)** control biofilm-trophic phage PaStL1. Grey border indicates strains overexpressing *sadC.* Data is representative of three independent experiments. Biofilm production of strains are indicated in **Supplementary Figures 2 and 3**.

Since glucose is also a subunit of Psl, it is unclear whether Bowman cannot infect these strains due to a direct requirement for LPS components or an indirect effect in which loss of *galU* and *ssg* disrupts glucose incorporation into Psl and impairs biofilm composition. The absence of biofilm-specific mutations in these PRMs suggests that Bowman may require the LPS core for infection, potentially as a secondary receptor.

Bowman was initially isolated alongside two other phages with incomplete genomes and low coverage. One of these phages had 99% identity to phage Pa5oct. This phage is reported to promote biofilm formation through *psl* upregulation and uses T4P, LPS, and possibly flagella for infection (91, 92). Pa5oct was not detectable in genomic sequencing after three rounds of plaque purification of Bowman but remained visible at low abundance by TEM (**Supplementary Figure S1**). This suggests a symbiotic relationship between the two phages, where Pa5oct promotes biofilm formation for Bowman to infect effectively (91, 92). Therefore, it may not be trivial to isolate Bowman alone. Pa5oct cannot infect PAO1 *fliC/pilA* or OSA-deficient strains but our hyper-biofilm strains lacking LPS O-antigens, T4P and/or flagella remained susceptible to the Bowman containing lysate (**Figure 5B**). Thus, the susceptibility data likely represent Bowman rather than Pa5oct infectivity.

### Lysates containing Bowman can kill carbapenem-resistant clinical isolates

Carbapenems are a powerful antibiotic class for treatment of recalcitrant *P. aeruginosa* infections. Since phage therapy is typically used as a last-resort approach for treatment of antibiotic-resistant infections, we next evaluated if Bowman could eradicate carbapenem-resistant clinical isolates. Growth of two (AR0094 and AR0095) of four clinical isolates tested was suppressed in the presence of Bowman **(Figure 7).** The two isolates are diverse, as indicated by differing MLST numbers (AR0094: ST115, AR0095: ST620) and have varying pilus activities (AR0094 twitches while AR0095 does not). Neither isolate formed significantly more biofilms than WT PAO1 (**Supplementary Figure S3**). This shows that Bowman may have clinical applicability in treatment of multidrug-resistant *P. aeruginosa* infections.

**Figure 7.**
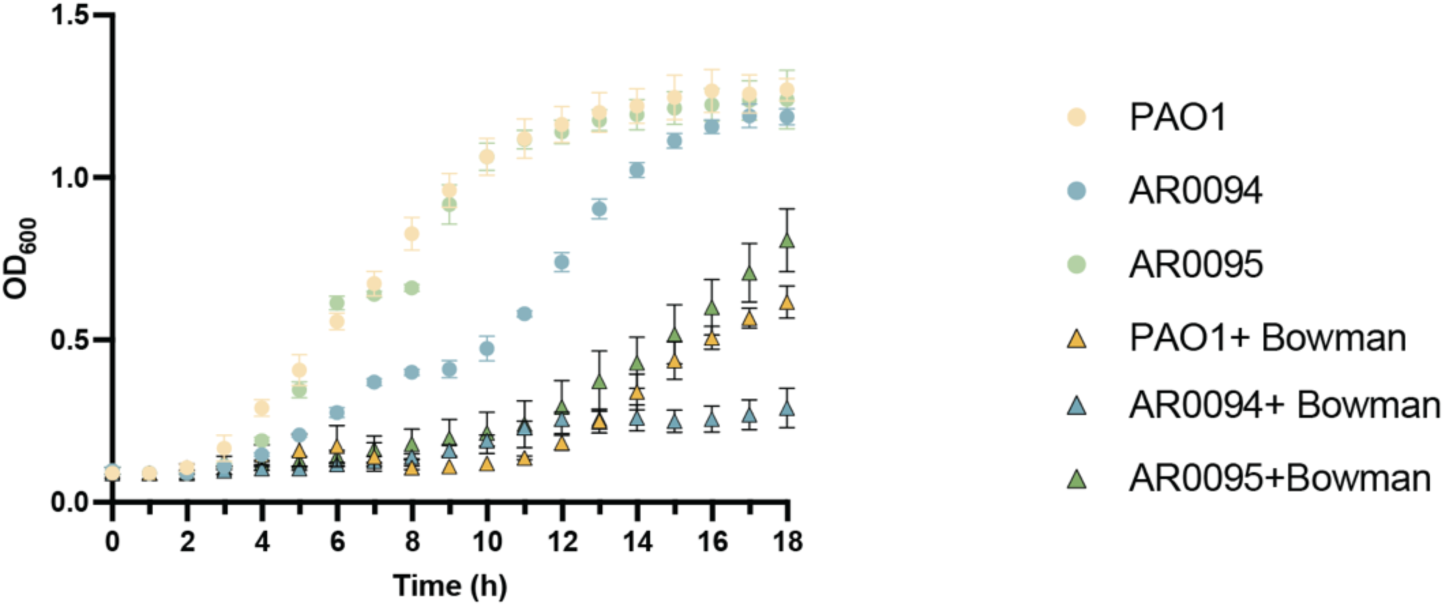
Bowman can kill certain carbapenem-resistant clinical isolates. Bowman killed two of the tested carbapenem resistant clinical isolates of different MLSTs. Data is representative of three independent experiments.

## DISCUSSION

Renewed interest in phages and advancements in sequencing technology have contributed to more rapid phage identification and characterization. Formal standards for naming, classification, and identification have not yet been widely established among bacteriophage researchers, unlike other areas of biology. Phage receptor usage, despite its key role in infection, is not consistently reported in the literature, likely due to the essential roles of some receptors and/or lack of bacterial strains that allow for ease of screening.

We therefore developed SURFAR to begin addressing this gap, as a defined and expandable set of strains in which receptor usage can be determined at the point of isolation. Many clinical strains encode multiple, complex anti-phage strategies but since the primary purpose of SURFAR is receptor characterization, such strains were not included in the platform. Instead, most strains are in the *P. aeruginosa* PAO1 background, which lacks these downstream anti-phage strategies, ensuring that resistance to phages likely due to receptor-based mechanisms. Further iterations of SURFAR, or SURFAR-like platforms may consider factors beyond receptor availability and incorporate mutants in additional downstream resistance strategies.

### Limitations, rationale, and expansion of SURFAR strains

One limitation of using PAO1 as the primary background is that the current version of SURFAR does not include all the 20 known O-antigen serotypes. PAO1 is serotype O5 and there are four other serotypes similar to O5: O2, O16, O18, and O20 (16). Resistance of our OSA mutants reflects an inability of the test phages to infect serotype O5 rather than general OSA-independence, as all our OSA mutants were O5-derived. Therefore, these phages may still require OSA of a different serogroup. Strains expressing other O-antigen serotypes, such as the other WT strains we have included, and their OSA counterparts would be useful additions to future versions of SURFAR.

To generate the overexpression strains, genes encoding for potential receptors were inserted into an arabinose inducible plasmid for titratable expression *in trans.* For phage isolation, specific receptor stoichiometry is likely not critical, as the goal is to select phages that can bind that receptor. Since phage adsorption is stochastic, increased receptor abundance could improve the probability that a phage successfully interacts with its target receptor (93). Alternative strategies such as elevating receptor expression through nutrient limitation or chromosomal mutations that constitutively inactivate repressors or activate regulators of receptor expression, may serve as alternate overexpression strategies. This iteration of SURFAR focused on TBDTs but SURFAR can be adapted to investigate other classes of potential receptors. For example, the porins and efflux pumps mutated in the ‘loss of potential receptors’ category could also be overexpressed in both WT and strains lacking common receptors T4P, LPS, and flagella to increase the probability of identifying phages that bind them.

To generate a set of hyper-biofilm producing strains, *sadC* was expressed *in trans*. This approach differs from other studies that increased biofilm production by deleting *wspF*, the methylesterase of the Wsp chemosensory system, to raise levels of intracellular cyclic-di-GMP (80, 81, 94). SadC is a *P. aeruginosa* digualylate cyclase (DCG) that is directly involved in cyclic-di-GMP synthesis and positively regulates Psl production. Compared to *wspR* (the DGC of the Wsp system) overexpression, *sadC* overexpression more effectively increases Psl synthesis (74, 95). Given the direct role of *sadC* in both Psl regulation and cyclic-di-GMP production, we selected this strategy as an alternative means of increasing biofilm production. In all of our strains, we observed increased biofilm production (**Supplementary Figure S2**) and biofilm-dependent phages more efficiently infected the overexpression strains (**Figure 5B**) suggesting this strategy was successful.

### Future expansion of SURFAR reference phages

To validate SURFAR, we tested its susceptibility to six distinct phages with known receptors. These reference phages span three receptor types (LPS core and O-antigen, pilus, and flagella). Most phages failed to infect the mutants lacking their required receptor but retained infectivity against other strains of the panel (**Figure 1**), confirming the expected patterns. When assessing Dalewood susceptibility, it failed to infect some strains carrying mutations in genes other than LPS core-related genes. These strains may encode additional undetected mutations, or the affected proteins may be required for infection as secondary receptors or co-factors. Ideally all strains included in SURFAR would be sequenced to rule out inclusion of spurious indels that could complicate interpretation.

Several SURFAR strains carry deletions in outer membrane proteins or their regulators, or overexpress outer membrane proteins. These mutations could indirectly alter outer membrane stability and structure. With the exceptions noted above, phages retained infectivity against strains expressing their cognate receptor, regardless of other mutations, indicating that such off-target effects are unlikely to affect phage infection.

As additional *P. aeruginosa* receptors are characterized and we find phages that do not depend on one or more of the common receptor types, the reference phage panel should be expanded. For example, biofilm-targeting phages such as Bowman and phages that use outer membrane receptors would be useful additions and comparators for phages with similar targets.

### Analysis of SURFAR isolated phage contigs

We showed that SURFAR can also be leveraged for phage isolation from wastewater samples. Of the 367 potential phage genomes and partial genomes identified, many could not be assigned to a known taxon and lacked sequence homology to entries in the NCBI BLAST and INPHARED databases. Contigs predicted to belong to the same taxon and/or sharing sequence similarity were often isolated by different strains, suggesting that including diverse strains allows SURFAR to enrich for diverse phages. Phage adsorption to a receptor is a random event as phages are not motile (93). Given this broad enrichment pattern, it remains unclear whether these probable phages productively infect *P. aeruginosa* or just adhere non-specifically. A key limitation of this study is that it represents a single replicate due to resource constraints. Further replicates that include wastewater samples from different sites and collection times are needed to more definitively correlate enrichment strains with resulting phage populations.

Approximately half of the phage contigs lacked informative BLAST hits, were orphans, and/or represented incomplete genomes. As a result, we cannot confidently determine if these contigs represent parts of complete phage genomes, non-specific extracellular genetic elements, or artifacts of sample preparation that include fragmented DNA. Since we sequenced enrichment lysates without additional purification or selection, our samples provide a crude snapshot of what is present. There were many contigs below the recommended cut off of 25x coverage (96), suggesting that our sequencing detected low abundance contigs. Environmental samples contain diverse content that may go undetected using traditional genomic sequencing techniques, often due to alternative or non-canonical nucleic acid compositions that phages use to evade host detection (97). As a result, these sequences can only be resolved at the class level (Caudoviricetes), or are not deposited into databases such as NCBI Virus or INPHARED and considered as novel or unknown. More advanced approaches, such as Oxford Nanopore and PacBio long read sequencing (98, 99), may detect these modifications and reduce the number of unclassified contigs.

Amongst the phage genomes with identifiable taxonomic homologs (**Figure 2A**), some come from phages may not target *P. aeruginosa*. *Felixounavirus, Klausavirus, Menderavirus, Tequintavirus, Tulanevirus, Veravirus*, and *Warwickvirus* phages have all been described to infect non-*Pseudomonas* genera (100–104). Phages enriched using a given strain are not limited to those capable of productive infection (e.g. infect and produce progeny or lysogenize), as they may also include phages that interact non-specifically with the enriching bacteria or were simply present at very high abundance in the starting filtrate. Since we did not perform subsequent rounds of sample purification, some of these non-specific phages may have remained in the samples at levels high enough for detection. An improvement to the workflow might include sequencing of wastewater filtrates prior to enrichment, to provide a reference snapshot of those phage genomes that were present. While such phages may not have direct clinical application for treatment of *P. aeruginosa* infections, their presence is still informative as it suggests their host bacteria co-exist with *P. aeruginosa* in the environment.

### Isolation and characterization of phages using SURFAR strains

Using SURFAR, we isolated and characterized the host range of two phages, Dalewood and Bowman, that are new members of previously described families. Based on host range, Dalewood appeared the LPS outer core as its receptor. Since the LPS outer core is more conserved than the O-antigen, a broader range of strains may be susceptible to Dalewood, consistent with its ability to infect multiple *P. aeruginosa* strains. Dalewood may also be combined with phages that use the LPS O-antigen, such as D3, as resistance to these phages often result in loss of O-antigen and exposure of the LPS outer core (105–107).

Bowman was unable to infect PAO1 *galU* or *ssg* mutants that overexpress the diguanylate cyclase SadC (**Figure 6A)**. PaStL1 also does not impair growth of PAO1 *galU* + pB-*sadC* or PAO1 *ssg* + pB-*sadC* **(Figure B)**. Products of *galU* and substrate that *ssg* acts on, is shared between Psl and the LPS outer core (28, 108). Therefore, it remains unclear whether these mutations cause resistance due to Bowman’s requirement for an intact LPS outer-core for infection or because of Psl disruption stemming from loss of a precursor. Other biofilm-targeting phages have not been shown to depend on LPS components for infection, making this potential receptor usage or link between the LPS outer-core and biofilms a novel finding. *wapH* is predicted to be involved in the addition of Glc II to the LPS core (28). Because this mutation likely affects only LPS structure and was one of the three loci mutated in a Bowman PRM, a clean *wapH* mutant may help reveal whether Bowman directly requires LPS core structures for infection (28). More PRMs may also be isolated because if Psl or other biofilm components are the primary receptor, mutations in genes of that pathway may be selected for.

### Isolation of Bowman and other biofilm-tropic phages

In some cases, a phage present in an enrichment lysate could not infect the strain initially used for enrichment. For example, Bowman was initially identified in a lysate enriched with WT PA1244 but did not form plaques on that strain and was instead propagated using PAO1 + pB*-sadC.* Since Bowman was isolated alongside Pa5oct, its initial ability to replicate using PA1244 may be the result of cooperation between the two phages, as Pa5Oct resistance is often accompanied by increased biofilm production (92). Bowman could not be isolated without trace amounts of Pa5oct (**Supplementary Figure S1**). This has implications for therapeutic application as a pure stock of Bowman may be difficult to prepare, and any contribution of Pa5oct would need to be accounted for. Entry into phage banks may require a pure phage but surprisingly, preparation to GMP standards does not explicitly state the same requirement. Instead, it generally requires preparations to be free from endotoxins and other residuals that may cause adverse effects (109, 110). Therefore, Bowman may have to be defined as a phage cocktail containing both phages, in which the ratio of both phages must be known. If pure Bowman is required, Bowman may be expressed and purified using its sequence, similar to protein purification strategies, in a Pa5oct-free condition.

Biofilm-tropic phages have previously been isolated using mutant strains that lack canonical receptors (80, 81). Our ability to isolate Bowman using a WT strain suggests that mutant strains may not be the only strategy. In fact, we isolated no phages of the *Bruynoghevirus* genus using strains with deletions of known receptors. Isolation of Bowman in combination with Pa5oct suggests that using an adjuvant phage or compound that can upregulate a potential receptor may be a promising technique for isolating phages that recognize new receptors. This mirrors receptor-based phage steering strategies in phage therapy where phages are administered to steer resistant bacteria to a phenotype relevant for infection, or in this case-phage isolation. For example, pre-treating enrichment hosts with Pa5oct selects for PRMs with elevated biofilm production-the condition that *Bruynoghevirus* requires for replication. Resistance to the first phage is a key step as it selects for bacterial phenotypes required to recover phages that recognize receptors that may be inaccessible or not typically expressed on wild-type cells.

Prior to the discovery of biofilm-tropic phages such as Bowman, biofilms were considered a barrier to phage efficacy as most phages cannot effectively penetrate biofilm structures to reach their cell-surface receptors. Biofilm-targeting phages address this barrier as they can reduce biofilm masses (80, 81). Further, biofilm-targeting phages may encode exopolysaccharide depolymerases that could be independently valuable. These depolymerases may be engineered independently of phages and used as therapeutic adjuvants in phage or antibiotic therapy. Using proteins or peptides rather than whole phages offers an additional benefit of bypassing the time-consuming process of phage isolation, purification, and cocktail preparation.

### Lack of Pel- and TBDT-targeting *P. aeruginosa* phages

Notably, no *P. aeruginosa* phages identified to date replicate in a Pel-dependent manner. In laboratory strain PAO1, Psl is the major exopolysaccharide in biofilms but in PA14, Pel is more abundant (111). The positive charge of Pel and its lower abundance in PAO1, the strain also used to isolate other biofilm-tropic phages, may explain the lack of Pel-targeting phages (111). It is unclear if Bowman requires Psl or Pel to infect, as *sadC* increases both.

*P. aeruginosa* phages that use TBDTs have not yet been identified. Unlike the three common receptors, TBDTs are present at lower surface density, strongly induced under iron limitation, and repressed when iron is available (46, 50, 112). Iron is one of the most abundant metals in wastewater (113) and standard enrichment media are iron-replete. Therefore, TBDTs may be expressed at low levels during sampling and enrichment. Phages that depend on these receptors may be poorly amplified in these conditions and outcompeted by phages that target constitutively expressed receptors, leaving them under-represented in isolated populations.

## Conclusion

As the incidence of antibiotic resistance rises and phage therapy improves, increasingly diverse *P. aeruginosa* phages have to be characterized and identified. Here, we describe and use a novel surface receptor-based platform for the isolation and characterization of novel *P. aeruginosa* phages. This iteration of SURFAR features overexpression and deletion of TonB-dependent receptors but the framework can be adapted to include additional mutants or strains that overexpress other classes of potential outer membrane protein receptors. Tools such as SURFAR are valuable for phage researchers as they help standardize characterization techniques and criteria. The adaptability of SURFAR to different strains, conditions, and media, and its user-friendly 96-well plate format makes it readily integrated by most labs.

## METHODS

### Bacterial strains and growth conditions

All strains and plasmids used in this study are outlined in Supplementary Table S2. Bacterial strains were grown in lysogeny broth (LB) at 37°C with shaking (200 rpm) or on 1.5% agar plates supplemented with antibiotics with the following concentrations, when required: gentamicin (Gm) 30 μg/mL, Gm 15 μg/mL for *E. coli*. L-arabinose was added to media at a concentration of 0.2% (w/v), when required, to induce expression from the pBADGr Ara promoter.

Deletion mutants were generated as previously described (49). Plasmid and chromosomal sequences were confirmed using Sanger sequencing (McMaster Genomics Facility) or Oxford Nanopore whole-plasmid sequencing (Plasmidsaurus). Primers are noted in Supplementary Table S2.

### Enrichment of wastewater with SURFAR strains

Wastewater effluents were filtered using 45μm syringe filters. Filtrates from four different sample sites were combined to a total volume of 40 mL to form the master filtered wastewater (MFW). One mL of LB was aliquoted into wells of a 96 deep well plate and 500 μL of MFW were added to each of the wells. Glycerol stocks of each of the SURFAR strains were inoculated into each of the wells. The whole plate was incubated at 37°C with shaking (200 rpm) overnight. Bacteria were then pelleted via centrifugation (2187 x g, 10 mins) and 1.5 mL of supernatant containing phages were carefully removed into individual Eppendorf tubes. 100 μL of chloroform was added to each tube to kill any remaining bacteria. These tubes were incubated at room temperature for 5 min, followed by high-speed vortex for 30 s, and incubation at room temperature for another 5 min. Tubes were centrifuged at 21,000 x *g* for 10 minutes to separate chloroform and lysates. Lysates were transferred to a new 1.5 mL Eppendorf tube and subjected to sequencing as described below.

### Phage genome sequencing and analysis

Phage lysate cultures were sequenced by the National Microbiology Lab in Winnipeg, MB. Lysates were treated with DNase I-XT (M0570, New England Biolabs) followed by genomic DNA extraction (Omega Bio-Tek). Short read libraries were prepared using the Nextera XT DNA library preparation kits and sequenced using the Illumina NextSeq 1000 or 2000 platform. FastQC v0.12.1 was used to assess quality of raw reads, trimmomatic v0.39 with a 4:20 sliding window was used to generate trimmed reads. Sequences were assembled using Spades (v4.0.0) and annotated using Pharokka (Version 1.7.3).

Contigs were assessed for homology using NCBI Virus BLAST and NCBI Microbes BLAST to generate Figure 2A.

Pairwise genomic distances between 367 sequenced phage contigs were estimated using MASH (v2.3) using default k-mer sketch patterns (21, 1000). All pairwise distances were computed using mash dist. The distance matrix was used to identify genome clusters using the NetworkX library (v.3.4.2). Sequence pairs sharing a MASH distance <0.05, sequence coverage >20x, and contig length >15,000 bp were connected as edges, connected components were defined as genome clusters. Results were visualized as a hierarchically clustered heatmap using seaborn (v0.13.2) and matplotlib (v.3.10.8). AI (Claude) was used to support the development of statistical code. Author has received the output and takes responsibility for the final content.

Bacteriophage genomes were extracted, sequenced, and assembled as previously described (17). Phages PO4 and Dalewood were amplified using *P. aeruginosa* PAO1 as the host strain, Bowman was amplified using *P. aeruginosa* PAO1 + pBADGR-*sadC*. Phage genomic DNA was extracted (Norgen Biotek Phage DNA Isolation Kit) and sequenced using the Illumina NextSeq 2000 platform (Microbial Genome Sequencing Center/SeqCenter, Pittsburgh, PA, USA). Sequencing results were assembled using SPAdes (version 3.15.5) (114) and annotated using Pharokka (version 1.7.3) (115).

The annotated genome of the phages are available at the following GenBank accession numbers: PO4-PX259653, Bowman-PZ584235, Dalewood-PZ594774.

### Transmission Electron Microscopy

Phages were amplified as described above to a titre of plaque forming unit (PFU)= 10^9^ and diluted 1:10. Phage lysate was applied to a 100kDA ultracentrifugal tube and centrifuged at 6076 x g for 5 min. The samples were washed with 5 mL of water, twice. One mL of sterile water was applied to collect the phage. Phage samples were submitted to the Canadian Centre for Electron Microscopy in Hamilton, ON for processing. Phages were imaged using a JEOL 1200EX microscope at 100,000X magnification.

### Selection of phage-resistant mutants

*P. aeruginosa* PAO1 was inoculated from overnight subculture into 5 mL LB and incubated at 37°C with shaking (200 rpm) to an optical density at 600 nm (OD600) of 0.6. Then, 100 μL of the bacterial culture was combined with 12 mL of 0.6% LB agar and 10 μL of PO4 at concentration of 10^8^ PFU in culture tubes and poured into petri dishes. Plates were incubated at 21°C for an additional 24 h or until the appearance of bacteriophage resistant colonies. Colonies were selected and grown on 1.5% LB agar, incubated at 37°C.

### Multi-strain phage susceptibility assays

Overnight cultures of bacteria were grown (1:1000) in liquid LB media and incubated at 37°C with shaking (200 rpm). Subcultures were standardized to OD600 of 0.5 in LB media. Phage stocks were standardized to 10^8^ PFU/mL and 10-fold serially diluted in phage buffer (68mM NaCl, 10mM Tris-HCl pH7.5, 10mM CaCl2, and 10mM MgSO4). Five μL each of culture and phage were combined and spotted onto 0.6% LB agar plates and dried for 5 min before incubation for 18-24 hours at 21°C. Phage susceptibility was indicated by absence of cell growth, relative to the control.

### Single strain phage plaquing assays

Overnight cultures of bacteria were grown (1:100) in liquid LB media and incubated at 37°C with shaking (200 rpm). Subcultures were standardized to OD600 of 0.3 in LB media and 100 μL of diluted culture was mixed with 12 mL of 0.6% LB agar and poured into standard petri dishes. LB agar was air dried in a biosafety cabinet. Phage stocks were standardized to 10^8^ PFU/mL and serially diluted with phage buffer. Five μL of each dilution was spotted onto the plates and air dried. Plates were incubated for 18 hours at 30°C. Phage susceptibility was indicated by phage plaque clearance. Plaque forming dilutions (PFD) defines the dilution in which single plaques were visible.

### Liquid phage susceptibility assay

Strains of interest were combined with Bowman or PaStL1 at multiplicity of infection (MOI)=1 in 100 μL LB. Samples were grown with shaking (orbital) at 37°C for 18 hours in the Biotek Epoch Plate readers. OD600 measurements were taken every 30 minutes. OD600 measurements were graphed using Graphpad Prism version 10.2.0. Phage susceptibility is indicated by lack of growth.

### Whole genome sequencing and mutation analysis of PRMs

Genomic DNA of *P. aeruginosa* PAO1 WT and phage resistant strains were extracted using the Wizard Genomic DNA Purification Kit. PRMs and PAO1 WT were sequenced using the Illumina NextSeq 2000 platform (Microbial Genome Sequencing Center/SeqCenter, Pittsburgh, PA, USA). All samples had average sequencing coverage of at least 50 reads. breseq (versions 0.35.4, 0.36.1, or 0.37.1) was used to identify mutations by comparing mutants sequences to the reference WT PAO1 Burrows lab strain (116). To ensure variant calling results were reliable, only those with more than 50 reads were included in the analysis unless specified (117). When required, mutations were confirmed by PCR and DNA sequencing (Plasmidsaurus, Oxford Nanopore).

### Data Visualization

Graphs and heat maps were generated using GraphPad Prism (10.5.0). Clustering was performed and visualized using seaborn clustermap (v0.13.2).

### Use of AI

AI (Claude) was used to support the development of statistical code used for Figures 2A and 3. The authors received the output and take responsibility for the final content. Claude was also used in editing of the chapter, but not its generation.

## Supporting information

Tran et al Supplementary Figs and Tables

## Data Availability

Annotated phage genome sequences were submitted to Genbank with the following accension numbers: Dalewood-PZ594774, Bowman-PZ584235, and Macklin-QB263992. Supplementary File S1 is available via the Zenodo digital repository at https://doi.org/10.5281/zenodo.23477247

## Acknowledgements

We thank Drs. Michael Parcey and Steven Gayder of the PhageSTAR program and Public Health Agency of Canada at the National Microbiology Laboratory for their assistance in phage sequencing and helpful discussions; Dr. Courtney Reichhardt for providing PaStL1; Dr. Lynne Howell for providing strains to include in the SURFAR platform; Dr. Derek Chan and Naaya Shah for generation of strains included in the SURFAR platform; Dr. Gerry Wright for providing clinical isolate strains; Dominique Tertigas and members of the Burrows lab for helpful discussions. This work was supported by a grant from the Canadian Institutes of Health Research to LLB (PJT-169053). LLB held the Canada Research Chair in Microbe-Surface Interactions (CRC-2021-00103), VNT holds a PGS-D award from the Natural Sciences and Engineering Research Council of Canada, IQ holds a CGS-D award from the Natural Sciences and Engineering Research Council of Canada.

## References

1. M. Naghavi, et al., Global burden of bacterial antimicrobial resistance 1990–2021: a systematic analysis with forecasts to 2050. The Lancet 404, 1199–1226 (2024).

2. A. Dublanchet, E. Fruciano, [A short history of phage therapy]. Med. Mal. Infect. 38, 415–420 (2008).

3. M. Żaczek, B. Weber-Dąbrowska, R. Międzybrodzki, M. Łusiak-Szelachowska, A. Górski, Phage Therapy in Poland – a Centennial Journey to the First Ethically Approved Treatment Facility in Europe. Front. Microbiol. 11 (2020).

4. J.-P. Pirnay, D. De Vos, G. Verbeken, Clinical application of bacteriophages in Europe. Microbiol. Aust. 40, 8 (2019).

5. Y.-F. Tsao, et al., Phage Morons Play an Important Role in *Pseudomonas aeruginosa* Phenotypes. J. Bacteriol. 200, 10.1128/jb.00189-18 (2018).

6. D. Holtappels, P. Alfenas-Zerbini, B. Koskella, Drivers and consequences of bacteriophage host range. FEMS Microbiol. Rev. 47, fuad038 (2023).

7. H. Harvey, et al., *Pseudomonas aeruginosa* defends against phages through type IV pilus glycosylation. Nat. Microbiol. 3, 47–52 (2018).

8. V. N. Tran, H. Harvey, T. S. Lahane, L. L. Burrows, The repertoire of resistance mutations selected by a *Pseudomonas aeruginosa* type IV pilus-targeting lytic bacteriophage. mBio 17, e00243–26 (2026).

9. V. N. Tran, L. L. Burrows, Steering the course: targeting and exploiting surface receptors in phage therapy. J. Bacteriol. 208, e00375–25 (2025).

10. J. M. Borin, et al., Rapid bacteria-phage coevolution drives the emergence of multiscale networks. Science 382, 674–678 (2023).

11. S. J. Labrie, J. E. Samson, S. Moineau, Bacteriophage resistance mechanisms. Nat. Rev. Microbiol. 8, 317–327 (2010).

12. T. Kaneko, K. Nakatsuka, S. Tsuneda, Phage cocktails: state-of-the-art technologies and strategies for effective design. FEMS Microbiol. Rev. 50, fuaf061 (2026).

13. D. Gao, et al., Fitness Trade-Offs in Phage Cocktail-Resistant *Salmonella enterica* Serovar Enteritidis Results in Increased Antibiotic Susceptibility and Reduced Virulence. Microbiol. Spectr. 10, e02914–22.

14. D. M. P. De Oliveira, et al., Antimicrobial Resistance in ESKAPE Pathogens. Clin. Microbiol. Rev. 33, e00181–19 (2020).

15. World Health Organization, WHO Bacterial Priority Pathogens List 2024: Bacterial Pathogens of Public Health Importance, to Guide Research, Development, and Strategies to Prevent and Control Antimicrobial Resistance, 1st ed (World Health Organization, 2024).

16. J. S. Lam, V. L. Taylor, S. T. Islam, Y. Hao, D. Kocíncová, Genetic and Functional Diversity of *Pseudomonas aeruginosa* Lipopolysaccharide. Front. Microbiol. 2, 118 (2011).

17. I. Qaderi, I. Chan, H. Harvey, L. L. Burrows, Structural conservation and functional role of TfpY-like proteins in type IV pilus assembly. J. Bacteriol. 0, e00343–24 (2025).

18. G. J. Newton, et al., Three-component-mediated serotype conversion in *Pseudomonas aeruginosa* by bacteriophage D3. Mol. Microbiol. 39, 1237–1247 (2001).

19. S. K. Arora, M. C. Wolfgang, S. Lory, R. Ramphal, Sequence Polymorphism in the Glycosylation Island and Flagellins of *Pseudomonas aeruginosa*. J. Bacteriol. 186, 2115–2122 (2004).

20. S. K. Arora, M. Bangera, S. Lory, R. Ramphal, A genomic island in *Pseudomonas aeruginosa* carries the determinants of flagellin glycosylation. Proc. Natl. Acad. Sci. 98, 9342–9347 (2001).

21. I. Qaderi, I. Chan, H. Harvey, L. L. Burrows, Structural conservation and functional role of TfpY-like proteins in type IV pilus assembly. J. Bacteriol. 0, e00343–24 (2025).

22. A. R. Costa, et al., Accumulation of defense systems in phage-resistant strains of*Pseudomonas aeruginosa*. Sci. Adv. 10, eadj0341 (2024).

23. J. A. Delgado-Nungaray, L. J. Figueroa-Yáñez, E. Reynaga-Delgado, A. M. Corona-España, O. Gonzalez-Reynoso, Unveiling the endogenous CRISPR-Cas system in *Pseudomonas aeruginosa* PAO1. PLOS ONE 19, e0312783 (2024).

24. C. L. Giltner, N. Rana, M. N. Lunardo, A. Q. Hussain, L. L. Burrows, Evolutionary and functional diversity of the *Pseudomonas* type IVa pilin island. Environ. Microbiol. 13, 250–264 (2011).

25. N. C. Esteves, D. N. Bigham, B. E. Scharf, Phages on filaments: A genetic screen elucidates the complex interactions between *Salmonella enterica* flagellin and bacteriophage Chi. PLOS Pathog. 19, e1011537 (2023).

26. J. S. Lam, H. L. Rocchetta, L. L. Burrows, Glycosyltransferases of *Pseudomonas aeruginosa* that assemble the O antigens of A band and B band lipopolysaccharide. J. Endotoxin Res. 5, 96–101 (1999).

27. H. L. Rocchetta, J. S. Lam, Identification and functional characterization of an ABC transport system involved in polysaccharide export of A-band lipopolysaccharide in *Pseudomonas aeruginosa*. J. Bacteriol. 179, 4713–4724 (1997).

28. J. D. King, D. Kocíncová, E. L. Westman, J. S. Lam, Review: Lipopolysaccharide biosynthesis in *Pseudomonas aeruginosa*. Innate Immun. 15, 261–312 (2009).

29. W. Rabsch, et al., FepA- and TonB-dependent bacteriophage H8: receptor binding and genomic sequence. J. Bacteriol. 189, 5658–5674 (2007).

30. B. K. Chan, et al., Phage selection restores antibiotic sensitivity in MDR *Pseudomonas aeruginosa*. Sci. Rep. 6, 26717 (2016).

31. L. N. Yaeger, et al., A genetic screen identifies a role for oprF in *Pseudomonas aeruginosa* biofilm stimulation by subinhibitory antibiotics. Npj Biofilms Microbiomes 10, 1–12 (2024).

32. S. Chevalier, et al., Structure, function and regulation of *Pseudomonas aeruginosa* porins. FEMS Microbiol. Rev. 41, 698–722 (2017).

33. E. G. Rawling, F. S. Brinkman, R. E. Hancock, Roles of the carboxy-terminal half of *Pseudomonas aeruginosa* major outer membrane protein OprF in cell shape, growth in low-osmolarity medium, and peptidoglycan association. J. Bacteriol. 180, 3556–3562 (1998).

34. E. K. Cassin, B. S. Tseng, Pushing beyond the Envelope: the Potential Roles of OprF in *Pseudomonas aeruginosa* Biofilm Formation and Pathogenicity. J. Bacteriol. 201, 10.1128/jb.00050-19 (2019).

35. X. Z. Li, H. Nikaido, K. Poole, Role of *mexA-mexB-oprM* in antibiotic efflux in *Pseudomonas aeruginosa*. Antimicrob. Agents Chemother. 39, 1948–1953 (1995).

36. T. Mine, Y. Morita, A. Kataoka, T. Mizushima, T. Tsuchiya, Expression in *Escherichia coli* of a new multidrug efflux pump, MexXY, from *Pseudomonas aeruginosa*. Antimicrob. Agents Chemother. 43, 415–417 (1999).

37. K. Poole, et al., Overexpression of the mexC-mexD-oprJ efflux operon in *nfxB*-type multidrug-resistant strains of *Pseudomonas aeruginosa*. Mol. Microbiol. 21, 713–724 (1996).

38. K. Jeannot, et al., Resistance and virulence of *Pseudomonas aeruginosa* clinical strains overproducing the MexCD-OprJ efflux pump. Antimicrob. Agents Chemother. 52, 2455–2462 (2008).

39. T. Köhler, S. F. Epp, L. K. Curty, J. C. Pechère, Characterization of MexT, the regulator of the MexE-MexF-OprN multidrug efflux system of *Pseudomonas aeruginosa*. J. Bacteriol. 181, 6300–6305 (1999).

40. P. D. Lister, D. J. Wolter, N. D. Hanson, Antibacterial-Resistant *Pseudomonas aeruginosa*: Clinical Impact and Complex Regulation of Chromosomally Encoded Resistance Mechanisms. Clin. Microbiol. Rev. 22, 582–610 (2009).

41. M. L. Sobel, S. Neshat, K. Poole, Mutations in PA2491 (*mexS*) Promote MexT-Dependent *mexEF-oprN* Expression and Multidrug Resistance in a Clinical Strain of *Pseudomonas aeruginosa*. J. Bacteriol. 187, 1246–1253 (2005).

42. P. E. Klebba, et al., Iron Acquisition Systems of Gram-negative Bacterial Pathogens Define TonB-Dependent Pathways to Novel Antibiotics. Chem. Rev. 121, 5193–5239 (2021).

43. N. Noinaj, M. Guillier, T. J. Barnard, S. K. Buchanan, TonB-dependent transporters: regulation, structure, and function. Annu. Rev. Microbiol. 64, 43–60 (2010).

44. L. Moynié, et al., The complex of ferric-enterobactin with its transporter from *Pseudomonas aeruginosa* suggests a two-site model. Nat. Commun. 10, 3673 (2019).

45. I. Josts, K. Veith, H. Tidow, Ternary structure of the outer membrane transporter FoxA with resolved signalling domain provides insights into TonB-mediated siderophore uptake. eLife 8, e48528 (2019).

46. D. C. K. Chan, L. L. Burrows, Pseudomonas aeruginosa FpvB Is a High-Affinity Transporter for Xenosiderophores Ferrichrome and Ferrioxamine B. mBio 14, e03149–22 (2022).

47. Q. Zhao, K. Poole, A second tonB gene in *Pseudomonas aeruginosa* is linked to the *exbB* and *exbD* genes. FEMS Microbiol. Lett. 184, 127–132 (2000).

48. M. R. M. Ranieri, et al., Thiostrepton Hijacks Pyoverdine Receptors To Inhibit Growth of *Pseudomonas aeruginosa*. Antimicrob. Agents Chemother. 63, e00472–19 (2019).

49. D. C. K. Chan, L. L. Burrows, Pseudomonas aeruginosa FpvB Is a High-Affinity Transporter for Xenosiderophores Ferrichrome and Ferrioxamine B. mBio 14, e03149–22 (2022).

50. N. Noinaj, M. Guillier, T. J. Barnard, S. K. Buchanan, TonB-dependent transporters: regulation, structure, and function. Annu. Rev. Microbiol. 64, 43–60 (2010).

51. M. A. Jacobs, et al., Comprehensive transposon mutant library of *Pseudomonas aeruginosa*. Proc. Natl. Acad. Sci. 100, 14339–14344 (2003).

52. L. R. Hmelo, et al., Precision-engineering the *Pseudomonas aeruginosa* genome with two-step allelic exchange. Nat. Protoc. 10, 1820–1841 (2015).

53. T. T. Hoang, R. R. Karkhoff-Schweizer, A. J. Kutchma, H. P. Schweizer, A broad-host-range Flp-FRT recombination system for site-specific excision of chromosomally-located DNA sequences: application for isolation of unmarked *Pseudomonas aeruginosa* mutants. Gene 212, 77–86 (1998).

54. M. L. Asikyan, J. V. Kus, L. L. Burrows, Novel Proteins That Modulate Type IV Pilus Retraction Dynamics in *Pseudomonas aeruginosa*. J. Bacteriol. 190, 7022–7034 (2008).

55. J. A. Moscoso, et al., The Diguanylate Cyclase SadC Is a Central Player in Gac/Rsm-Mediated Biofilm Formation in *Pseudomonas aeruginosa*. J. Bacteriol. 196, 4081–4088 (2014).

56. N. A. Roberge, L. L. Burrows, Building permits—control of type IV pilus assembly by PilB and its cofactors. J. Bacteriol. 206, e00359–24 (2024).

57. R. Cook, et al., Bacteriophage genomics: What has five years of INPHARED taught us? [Preprint] (2026). Available at: https://www.biorxiv.org/content/10.64898/2026.05.06.722914v1 [Accessed 6 August 2026].

58. D. E. Bradley, Basic Characterization of a *Pseudomonas aeruginosa* Pilus-Dependent Bacteriophage with a Long Noncontractile Tail. J. Virol. 12, 1139–1148 (1973).

59. A. M. Kropinski, Sequence of the Genome of the Temperate, Serotype-Converting, P*Pseudomonas aeruginosa* Bacteriophage D3. J. Bacteriol. **182**, 6066–6074 (2000).

60. I. Qaderi, et al., Structural diversification of phage tail fibers enables recognition of diverse type IV pili. Proc. Natl. Acad. Sci. 123, e2626959123 (2026).

61. A. Washizaki, T. Yonesaki, Y. Otsuka, Characterization of the interactions between *Escherichia coli* receptors, LPS and OmpC, and bacteriophage T4 long tail fibers. MicrobiologyOpen 5, 1003–1015 (2016).

62. K. E. Kortright, B. K. Chan, P. E. Turner, High-throughput discovery of phage receptors using transposon insertion sequencing of bacteria. Proc. Natl. Acad. Sci. U. S. A. 117, 18670–18679 (2020).

63. W.-X. Yee, et al., Intracellular barriers and receptor masking limit success of a Pseudomonas aeruginosa clinical phage family. [Preprint] (2026).

64. M. C. Johnson, et al., Core defense hotspots within *Pseudomonas aeruginosa* are a consistent and rich source of anti-phage defense systems. Nucleic Acids Res. 51, 4995–5005 (2023).

65. A. Verma, et al., Glycosylation of b-Type Flagellin of *Pseudomonas aeruginosa*: Structural and Genetic Basis. J. Bacteriol. 188, 4395–4403 (2006).

66. A. Jurczak-Kurek, et al., Biodiversity of bacteriophages: morphological and biological properties of a large group of phages isolated from urban sewage. Sci. Rep. 6, 34338 (2016).

67. N. M. Alharbi, M. M. Ziadi, Wastewater as a fertility source for novel bacteriophages against multi-drug resistant bacteria. Saudi J. Biol. Sci. 28, 4358–4364 (2021).

68. D. Turner, A. M. Kropinski, E. M. Adriaenssens, A Roadmap for Genome-Based Phage Taxonomy. Viruses 13, 506 (2021).

69. D. Amgarten, B. K. V. Iha, C. M. Piroupo, A. M. da Silva, J. C. Setubal, vHULK, a New Tool for Bacteriophage Host Prediction Based on Annotated Genomic Features and Neural Networks. PHAGE Ther. Appl. Res. 3, 204–212 (2022).

70. S. Roux, et al., iPHoP: An integrated machine learning framework to maximize host prediction for metagenome-derived viruses of archaea and bacteria. PLOS Biol. 21, e3002083 (2023).

71. D. a. C. Heesterbeek, et al., Complement-dependent outer membrane perturbation sensitizes Gram-negative bacteria to Gram-positive specific antibiotics. Sci. Rep. 9, 3074 (2019).

72. K. Danis-Wlodarczyk, et al., Characterization of the Newly Isolated Lytic Bacteriophages KTN6 and KT28 and Their Efficacy against *Pseudomonas aeruginosa* Biofilm. PLOS ONE 10, e0127603 (2015).

73. F. Li, et al., High-resolution cryo-EM structure of the *Pseudomonas* bacteriophage E217. Nat. Commun. 14, 4052 (2023).

74. X. Zhu, et al., A comparative analysis of phage classification methods in light of the recent ICTV taxonomic revisions. Virology 594, 110016 (2024).

75. A. Lopes, P. Tavares, M.-A. Petit, R. Guérois, S. Zinn-Justin, Automated classification of tailed bacteriophages according to their neck organization. BMC Genomics 15, 1027 (2014).

76. L. Girard, C. Lood, R. De Mot, V. van Noort, J. Baudart, Genomic diversity and metabolic potential of marine *Pseudomonadaceae*. Front. Microbiol. 14 (2023).

77. A. L. B. Canellas, M. S. Laport, Marine *Pseudomonas*: diving into the waves of blue biotechnology. Lett. Appl. Microbiol. 77, ovae099 (2024).

78. A. V. Nair, et al., A comparative study of coastal and clinical isolates of *Pseudomonas aeruginosa*. Braz. J. Microbiol. 46, 725–734 (2015).

79. L. Meneses, et al., Directed evolution of phages in biofilms enhances *Pseudomonas aeruginosa*control through improved lipopolysaccharide recognition. Nat. Commun. 16, 10219 (2025).

80. B. Walton, et al., A biofilm-tropic Pseudomonas aeruginosa bacteriophage uses the exopolysaccharide Psl as receptor. [Preprint] (2024). Available at: https://elifesciences.org/reviewed-preprints/102352v1 [Accessed 31 March 2025].

81. K. Amyx-Sherer, et al., Two unrelated *Pseudomonas aeruginosa* phages require the exopolysaccharide Psl for infection. Npj Biofilms Microbiomes 11, 211 (2025).

82. Z. Chegini, et al., Bacteriophage therapy against *Pseudomonas aeruginosa* biofilms: a review. Ann. Clin. Microbiol. Antimicrob. 19, 45 (2020).

83. L. Ma, H. Lu, A. Sprinkle, M. R. Parsek, D. J. Wozniak, *Pseudomonas aeruginosa* Psl Is a Galactose- and Mannose-Rich Exopolysaccharide. J. Bacteriol. 189, 8353–8356 (2007).

84. M. S. Byrd, B. Pang, M. Mishra, W. E. Swords, D. J. Wozniak, The *Pseudomonas aeruginosa* Exopolysaccharide Psl Facilitates Surface Adherence and NF-κB Activation in A549 Cells. mBio 1, 10.1128/mbio.00140-10 (2010).

85. D. Gutiérrez, et al., Role of the Pre-neck Appendage Protein (Dpo7) from Phage vB_SepiS-phiIPLA7 as an Anti-biofilm Agent in *Staphylococcal* Species. Front. Microbiol. 6 (2015).

86. Y. Wu, et al., A Novel Polysaccharide Depolymerase Encoded by the Phage SH-KP152226 Confers Specific Activity Against Multidrug-Resistant *Klebsiella pneumoniae* via Biofilm Degradation. Front. Microbiol. 10 (2019).

87. A. Cornelissen, et al., Identification of EPS-degrading activity within the tail spikes of the novel *Pseudomonas putida* phage AF. Virology 434, 251–256 (2012).

88. A. Cornelissen, et al., The T7-related *Pseudomonas putida* phage φ15 displays virion-associated biofilm degradation properties. PloS One 6, e18597 (2011).

89. S. Le, et al., Chromosomal DNA deletion confers phage resistance to *Pseudomonas aeruginosa*. Sci. Rep. 4, 4738 (2015).

90. C. R. Dean, J. B. Goldberg, *Pseudomonas aeruginosa galU* is required for a complete lipopolysaccharide core and repairs a secondary mutation in a PA103 (serogroup O11) *wbpM* mutant. FEMS Microbiol. Lett. 210, 277–283 (2002).

91. C. Lood, et al., Integrative omics analysis of *Pseudomonas aeruginosa* virus PA5oct highlights the molecular complexity of jumbo phages. Environ. Microbiol. 22, 2165–2181 (2020).

92. T. Olszak, et al., *Pseudomonas aeruginosa* PA5oct Jumbo Phage Impacts Planktonic and Biofilm Population and Reduces Its Host Virulence. Viruses 11, 1089 (2019).

93. S. T. Abedon, Bacteriophage Adsorption: Likelihood of Virion Encounter with Bacteria and Other Factors Affecting Rates. Antibiotics 12, 723 (2023).

94. J. W. Hickman, D. F. Tifrea, C. S. Harwood, A chemosensory system that regulates biofilm formation through modulation of cyclic diguanylate levels. Proc. Natl. Acad. Sci. 102, 14422–14427 (2005).

95. Y. Irie, et al., Self-produced exopolysaccharide is a signal that stimulates biofilm formation in *Pseudomonas aeruginosa*. Proc. Natl. Acad. Sci. U. S. A. 109, 20632–20636 (2012).

96. D. Turner, E. M. Adriaenssens, I. Tolstoy, A. M. Kropinski, Phage Annotation Guide: Guidelines for Assembly and High-Quality Annotation. Phage 2, 170–182 (2021).

97. A. R. Weinheimer, et al., Single-particle genomics uncovers abundant non-canonical marine viruses from nanolitre volumes. Nat. Microbiol. 10, 3245–3257 (2025).

98. L. K. White, J. R. Hesselberth, Modification mapping by nanopore sequencing. Front. Genet. 13 (2022).

99. T. K. Nielsen, et al., Detection of nucleotide modifications in bacteria and bacteriophages: Strengths and limitations of current technologies and software. Mol. Ecol. 32, 1236–1247 (2023).

100. C. L. Schoch, et al., NCBI Taxonomy: a comprehensive update on curation, resources and tools. Database 2020, baaa062 (2020).

101. R. Barron-Montenegro, et al., Comparative Analysis of Felixounavirus Genomes Including Two New Members of the Genus That Infect *Salmonella infantis*. Antibiotics 10, 806 (2021).

102. D. Damnjanović, X. Vázquez-Campos, L. Elliott, M. Willcox, W. J. Bridge, Characterisation of Bacteriophage vB_SmaM_Ps15 Infective to *Stenotrophomonas maltophilia* Clinical Ocular Isolates. Viruses 14, 709 (2022).

103. W. Liu, et al., Complete genome sequencing of a *Tequintavirus* bacteriophage with a broad host range against *Salmonella abortus equi* isolates from donkeys. Front. Microbiol. 13, 938616 (2022).

104. A. R. Vitt, M. C. H. Sørensen, V. Bortolaia, L. Brøndsted, A Representative Collection of Commensal Extended-Spectrum- and AmpC-β-Lactamase-Producing *Escherichia coli* of Animal Origin for Phage Sensitivity Studies. PHAGE 4, 35–45 (2023).

105. R. C. T. Wright, V.-P. Friman, M. C. M. Smith, M. A. Brockhurst, Cross-resistance is modular in bacteria-phage interactions. PLoS Biol. 16, e2006057 (2018).

106. Y. Yang, et al., Development of a Bacteriophage Cocktail to Constrain the Emergence of Phage-Resistant *Pseudomonas aeruginosa*. Front. Microbiol. 11, 327 (2020).

107. G. Li, et al., Adaptation of *Pseudomonas aeruginosa* to Phage PaP1 Predation via O-Antigen Polymerase Mutation. Front. Microbiol. 9 (2018).

108. M. S. Byrd, et al., Genetic and Biochemical Analyses of the *Pseudomonas aeruginosa* Psl Exopolysaccharide Reveal Overlapping Roles for Polysaccharide Synthesis Enzymes in Psl and LPS Production. Mol. Microbiol. 73, 622–638 (2009).

109. L. Bretaudeau, K. Tremblais, F. Aubrit, M. Meichenin, I. Arnaud, Good Manufacturing Practice (GMP) Compliance for Phage Therapy Medicinal Products. Front. Microbiol. 11 (2020).

110. M. Fuerst-Wilmes, et al., Regulation of phage therapy medicinal products: developments, challenges, and opportunities. Front. Cell. Infect. Microbiol. 15, 1631359 (2025).

111. K. M. Colvin, et al., The Pel and Psl polysaccharides provide *Pseudomonas aeruginosa* structural redundancy within the biofilm matrix. Environ. Microbiol. 14, 10.1111/j.1462-2920.2011.02657.x (2012).

112. P. E. Klebba, M. A. McIntosh, J. B. Neilands, Kinetics of biosynthesis of iron-regulated membrane proteins in *Escherichia coli*. J. Bacteriol. 149, 880–888 (1982).

113. E. Risch, L. Jaumaux, C. Maeseele, J.-M. Choubert, Comparative Life Cycle Assessment of two advanced treatment steps for wastewater micropollutants: How to determine whole-system environmental benefits? Sci. Total Environ. 805, 150300 (2022).

114. A. Prjibelski, D. Antipov, D. Meleshko, A. Lapidus, A. Korobeynikov, Using SPAdes De Novo Assembler. 10.1002/cpbi.102.

115. G. Bouras, et al., Pharokka: a fast scalable bacteriophage annotation tool. Bioinformatics 39, btac776 (2023).

116. D. E. Deatherage, J. E. Barrick, Identification of mutations in laboratory evolved microbes from next-generation sequencing data using breseq. Methods Mol. Biol. Clifton NJ 1151, 165–188 (2014).

117. A. Petrackova, et al., Standardization of Sequencing Coverage Depth in NGS: Recommendation for Detection of Clonal and Subclonal Mutations in Cancer Diagnostics. Front. Oncol. 9 (2019).

