## Supplementary material for "Identification and characterization of new *P. aeruginosa* phages using a surface receptor platform": Tran et al Supplementary Figs and Tables

Veronica N. Tran, Tanisha S. Lahane, Ikram Qaderi, Hanjeong Harvey and Lori L. Burrows*
Department of Biochemistry and Biomedical Sciences and the Michael G. DeGroote Institute for Infection Disease Research, McMaster University

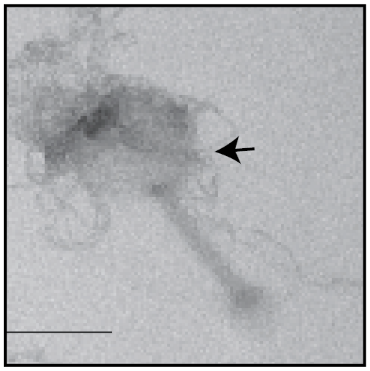

**Supplementary Figure S1. TEM image of Pa5oct contaminant in Bowman sample.** Arrow indicates phage capsid. Filaments are predicted to be biofilm exopolysaccharides. Scale bars = 100 nm.

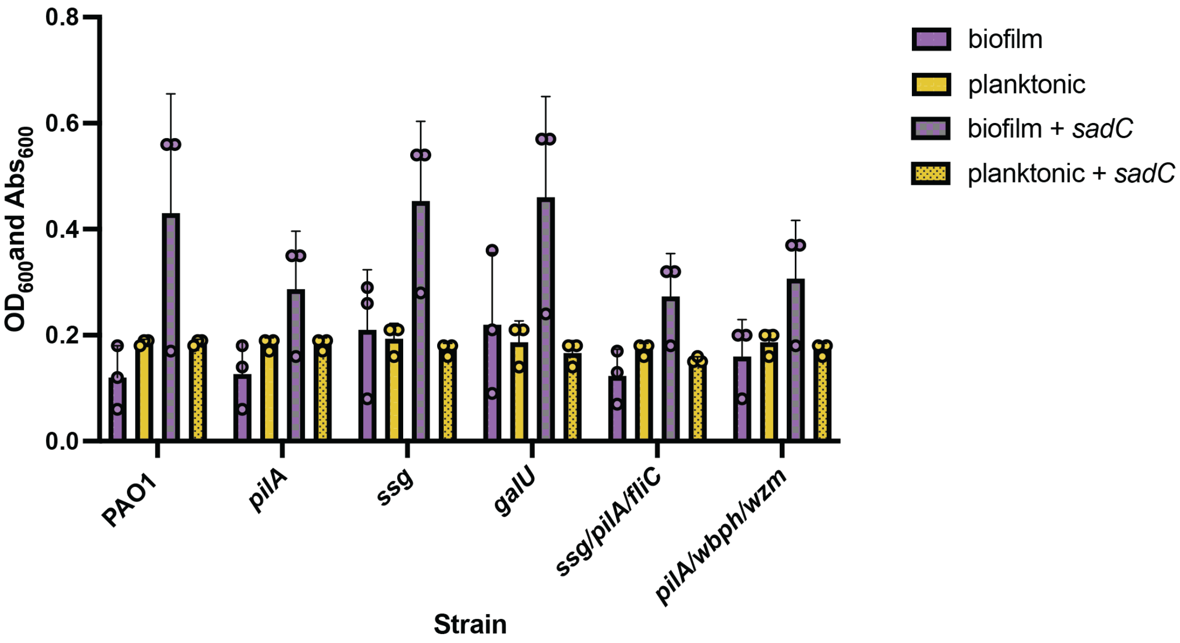


**Supplementary Figure S2. Biofilm and planktonic growth of LPS mutant strains from Figure 5A and B.** *sadC* overexpression strains produce more biofilms than vector controls. Data is representative of three independent experiments.

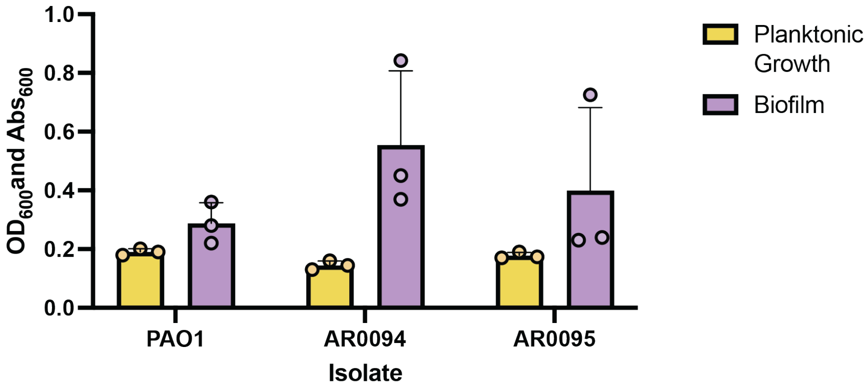


**Supplementary Figure S3. Biofilm and planktonic growth of clinical isolates from Figure 5C.** There are no statistical differences in biofilm formation between the two clinical isolates and PAO1. Data is representative of three independent experiments.

**Supplementary Table S1 Strains that enriched for each cluster in Figure 5B**

| Cluster | Strains |
|---|---|
| **Cluster 1** | PA1457 *pilA* |
| | PAO1 *fliC* |
| | PAK *fliC* |
| | PAO1 *fliC/pilA* |
| | PAO1 *ssg* |
| | PAO1 WT |
| | PAO1 *ssg/pilA/fliC* |
| | PAO1 + pB-*fpvB* |
| | PAO1 *pilA/wbpH/wzm* + pB-*foxA* |
| | PAO1 *pilA/wbpH/wzm* + pB-*piuA* |
| | PAO1 *pilA/wbpH/wzm* + pB-*fiuA* |
| | PAO1 *pilA/wbpH/wzm* + pB-*sadC* |
| | PAO1 fliC/*pilA* + pB-*sadC* |
| | PA14 *pilA* |
| **Cluster 2** | PA1457 *pilA* |
| | PAO1 *fliC* |
| | PAK *fliC* |
| | PAO1 *fliC/pilA* |
| | PAO1 *ssg* |
| | PAO1 |
| | PAO1 *ssg/pilA/fliC* |
| | PAO1 + pB-*fpvB* |
| | PAO1 *pilA/wbpH/wzm* + pB-*piuA* |
| | PAO1 *pilA/wbpH/wzm* + pB-*fiuA* |
| | PAO1 *pilA/wbpH/wzm* + pB-*sadC* |
| | PAO1 fliC/*pilA* + pB-*sadC* |
| | PA14 *pilA* |
| **Cluster 3** | PAO1 *fliC* |
| | PAO1 *galU* |
| | PAO1 |
| | PAO1 *tonB2 (1)* |
| | PAO1 *tonB2 (2)* |
| | PAO1 *pelA* |
| | PAO1 *psl (1)* |
| | PAO1 *psl (2)* |
| | PAO1 + pB-fiuA |

| | |
|---|---|
| **Cluster 4** | PAO1 *pilA/wbpH/wzm* |
| | PAO1 *tonB1 (1)* |
| | PAO1 *tonB1 (2)* |
| | PA14 |
| | PAO1 *mexT* |
| | PAO1 pilA |
| **Cluster 5** | PAO1 *wbpH* |
| | PAO1 *wzm* |
| | PAO1 *mexT* |
| | PAO1 + pB-*fpvA* |
| | PAO1 +pB-*foxA* |
| | PAO1 *pilA/wbpH/wzm* + pB-*fpvA* |
| **Cluster 6** | PA5196 |
| | PA5196 *pilA* |
| **Cluster 7** | PAO1 *fliC/pilA* |
| | PAO1 fliC/*pilA* + pB-*sadC* |
| **Cluster 8** | PAO1 *fliC/pilA* |
| | PAO1 fliC/*pilA* + pB-*sadC* |
| **Cluster 9** | PA1244 |
| | PA1244 *pilA* |
| **Cluster 10** | PA1244 |
| | PA1244 *pilA* |
| **Cluster 11** | PAO1 + pB-*fpvA* |
| | PAO1 + pB-*fpvB* |
| **Cluster 12** | PA1457 |
| | PA1244 |
| pB=pBADGr | |

**Supplementary Table S2. Strains and plasmids used in this study**

| **Strain/plasmid** | **Characteristics** | **Source** |
|---|---|---|
| *P. aeruginosa strains* | | |
| mPAO1 | WT, group II pilins | lab collection |
| mPAO1 *pilA::Tn5/NP* | Tn5-phoA insertion at position of 163 in *pilA* | (1) |
| PA1244 | Pilin Group I clinical isolate | (2) |
| PA14 | Group III pilin lab strain | (3) |
| PA5196 | Group IV pilin clinical isolate | (4) |
| PA1457 | Group V pilin clinical isolate | (5) |
| PA1244 NP | FRT insertion in *pilA* | lab collection |

| | | |
|---|---|---|
| PA14 NP | Deletion of *pilA* | (6) |
| PA5196 NP | *pilA::aacC1FRT* | (7) |
| PA1457 NP | FRT insertion in *pilA* | lab collection |
| mPAO1 Δ*fliC* | mPAO1 fliC deletion | lab collection |
| PAK *fliC* | FRT insertion in *fliC* | (8) |
| mPAO1 *flgE* | mPAO1 *flgE* deletion | lab collection |
| mPAO1 Δ*wbpH* | mPAO1 *wbpH* deletion | lab collection |
| mPAO1 Δ*wzm* | mPAO1 *wzm* deletion | lab collection |
| mPAO1 Δ*ssg* | mPAO1 *ssg* deletion | lab collection |
| mPAO1 Δ*galU* | mPAO1 *galU* deletion | lab collection |
| mPAO1 Δ*fliC/pilA pilA::Tn5/NP* | mPAO1 with *fliC* deletion and *pilA* transposon | lab collection |
| mPAO1 Δ*ssg*/Δ*pilA* | mPAO1 with *ssg* and *pilA* deletions | lab collection |
| mPAO1 Δ*ssg*/Δ*pilA*/Δ*fliC* | mPAO1 with *ssg, pilA,* and *fliC* deletions | lab collection |
| mPAO1 Δ*tonB1* | mPAO1 with *tonB1* deletion | this work |
| mPAO1 Δ*tonB2* | mPAO1 with *ton2* deletion | this work |
| mPAO1 Δ*fpvA* | mPAO1 with *fpvA* deletion | this work |
| mPAO1 Δ*fpvB* | mPAO1 with *fpvB* deletion | this work |
| mPAO1 Δ*fptA* | mPAO1 with *fptA* deletion | this work |
| mPAO1 *oprF::HiMar1* | Himar1 transposon insertion mutant in *oprF* *Libnrary ID: BBtn1_G3* | (9) |
| mPAO1 *oprJDM* | Deletion of *oprJDM* | lab collection |
| mPAO1 Δ*mexT* | mPAO1 with *mexT* deletion | this work |
| mPAO1 Δ*pelA* | PAO1 with *pelA* deletion | (10) |
| mPAO1 Δ*psl* | mPAO1 with *pslBCD* deletion | this work |
| mPAO1+pBADGr-*fpvA* | mPAO1 complemented with *fpvA* in pBADGr | this work |
| mPAO1+pBADGr-*fpvB* | mPAO1 complemented with *fpvB* in pBADGr | this work |
| mPAO1+pBADGr-*fptA* | mPAO1 complemented with *fptA* in pBADGr | this work |
| mPAO1+pBADGr-*foxA* | mPAO1 complemented with *foxA* in pBADGr | this work |
| mPAO1+pBADGr-*piuA* | mPAO1 complemented with *piuA* in pBADGr | this work |
| mPAO1+pBADGr-*fiuA* | mPAO1 complemented with *fiuA* in pBADGr | this work |
| mPAO1*pilA/wbpH/wzm* +pBADGr-*fpvA* | mPAO1 *pilA/wbpH/wzm* complemented with *fpvA* in pBADGr | this work |
| mPAO1*pilA/wbpH/wzm* +pBADGr-*fpvB* | mPAO1 *pilA/wbpH/wzm* complemented with *fpvB* in pBADGr | this work |

| | | |
|---|---|---|
| mPAO1*pilA/wbpH/wzm* +pBADGr-*fptA* | mPAO1 *pilA/wbpH/wzm* complemented with *fptA* in pBADGr | this work |
| mPAO1*pilA/wbpH/wzm* +pBADGr-*foxA* | mPAO1 *pilA/wbpH/wzm* complemented with *foxA* in pBADGr | this work |
| mPAO1*pilA/wbpH/wzm* +pBADGr-*piuA* | mPAO1 *pilA/wbpH/wzm* complemented with *piuA* in pBADGr | this work |
| mPAO1*pilA/wbpH/wzm* +pBADGr-*fiuA* | mPAO1 *pilA/wbpH/wzm* complemented with *fiuA* in pBADGr | this work |
| mPAO1 +pBADGr-*sadC* | mPAO1 complemented with *sadC* in pBADGr | (11) |
| mPAO1 *pilA/wbpH/wzm* +pBADGr-*sadC* | mPAO1 *pilA/wbpH/wzm* complemented with *sadC* in pBADGr | this work |
| mPAO1 *fliC/pilA* +pBADGr-*sadC* | mPAO1 *fliC/pilA* complemented with *sadC* in pBADGr | this work |
| mPAO1 *galU* +pBADGr-*sadC* | mPAO1 *galU* complemented with *sadC* in pBADGr | this work |
| mPAO1 *ssg* +pBADGr-*sadC* | mPAO1 *ssg* complemented with *sadC* in pBADGr | this work |
| *E. coli strains* | | |
| DH5α | *F-φ80lacZΔM15 Δ(lacZYA-argF)U169 recA1 endA1 hsdR17(rk−, mk+) phoA supE44 thi-1 gyrA96 relA1 λ−* | Invitrogen |
| SM10 | *thi-1 thr leu tonA lacY supE recA::RP4-2-Tc::Mu (KmR)* | Invitrogen |
| Plasmids | | |
| pEX18Gm | Suicide vector for gene replacement | (12) |
| pBADGr | arabinose-inducible, broad host range complementation vector | (13) |
| pBADGr-*sadC* | pBADGr expressing *sadC* | (11) |
| pBADGr-*fpvA* | pBADGr expressing *fpvA* | this work |
| pBADGr-*fpvB* | pBADGr expressing *fpvB* | this work |
| pBADGr-*fptA* | pBADGr expressing *fptA* | this work |
| pBADGr-*foxA* | pBADGr expressing *foxA* | this work |
| pBADGr-*piuA* | pBADGr expressing *piuA* | this work |
| pBADGr-*fiuA* | pBADGr expressing *fiuA* | this work |
| pEX18GM- *fpvA* | Construct to delete *fpvA* | (14) |
| pEX18GM- *fpvB* | Construct to delete *fpvB* | (14) |
| pEX18GM-*fptA* | Construct to delete *fptA* | (14) |
| Overexpression primers | | |
| Gene | Forward (5' -> 3') | Reverse (5' -> 3') |

| | | |
|---|---|---|
| *fpvA* | CAATTCTAGACGCAACTTGCCAACCTAAGAAGAG | CAATCTGCAGGGGCGCTGATCTCAGGTCCGTTCC |
| *fpvB* | CAATGAATTCTGCGCCGCCATCCAGGACACTGCA | CAATAAGCTTCGAGCGAGTTCGAGCGCCGCTATC |
| *foxA* | CAATTCTAGATTCAACGACTCCAACGGATTCC | CAGTAAGCTTGTAGCCAGACCGACATAGCG |
| *fiuA* | AATGAATTCCATCCAGGAGCCCCGCACATG | AATAAGCTTTCACCACTTGTAGCTGACGCTG |
| *fptA* | ACAACCCGGGGAGGCTGTTCGAACGATGAAAACG | AACAAAGCTTTCAGAACGCGCCCCGC |
| *piuA* | ATAAGAATTCATTCCGCGACCTTCACAGCTCC | CCAAGCTTGCCTTGTCGCTGTCCATCTCG |

### Supplementary References


1. M. A. Jacobs, *et al.*, Comprehensive transposon mutant library of *Pseudomonas aeruginosa*. *Proceedings of the National Academy of Sciences* **100**, 14339–14344 (2003).
2. P. A. Castric, C. D. Deal, Differentiation of *Pseudomonas aeruginosa* pili based on sequence and B-cell epitope analyses. *Infect Immun* **62**, 371–376 (1994).
3. L. G. Rahme, *et al.*, Common virulence factors for bacterial pathogenicity in plants and animals. *Science* **268**, 1899–1902 (1995).
4. J. Kus, D. Tullis, D. Cvitkovitch, L. Burrows, Significant differences in type IV pilin allele distribution among *Pseudomonas aeruginosa* isolates from cystic fibrosis (CF) versus non-CF patients. *Microbiology (Reading, England)* **150**, 1315–26 (2004).
5. R. Barnshaw, *et al.*, Separation of Pseudomonas aeruginosa type IV pilus-dependent twitching motility and surface-sensing responses. *mBio* **16**, e02521-25 (2025).
6. Y. Nguyen, *et al.*, A Highly Dynamic Loop of the *Pseudomonas aeruginosa* PA14 Type IV Pilin Is Essential for Pilus Assembly. *ACS Infect. Dis.* **4**, 936–943 (2018).
7. J. V. Kus, *et al.*, Modification of P*seudomonas aeruginosa* Pa5196 Type IV Pilins at Multiple Sites with d-Araf by a Novel GT-C Family Arabinosyltransferase, TfpW. *Journal of Bacteriology* **190**, 7464–7478 (2008).
8. S. L. N. Kilmury, L. L. Burrows, The *Pseudomonas aeruginosa* PilSR Two-Component System Regulates Both Twitching and Swimming Motilities. *mBio* **9**, e01310-18 (2018).
9. L. N. Yaeger, *et al.*, A genetic screen identifies a role for oprF in *Pseudomonas aeruginosa* biofilm stimulation by subinhibitory antibiotics. *npj Biofilms Microbiomes* **10**, 1–12 (2024).
10. M. T. Rybtke, *et al.*, Fluorescence-Based Reporter for Gauging Cyclic Di-GMP Levels in Pseudomonas aeruginosa. *Applied and Environmental Microbiology* **78**, 5060–5069 (2012).
11. N. Roberge, *et al.*, Twitching motility suppressors reveal a role for FimX in type IV pilus extension dynamics. *PLOS Genetics* **21**, e1011802 (2025).
12. T. T. Hoang, R. R. Karkhoff-Schweizer, A. J. Kutchma, H. P. Schweizer, A broad-host-range Flp-FRT recombination system for site-specific excision of chromosomally-located DNA sequences: application for isolation of unmarked *Pseudomonas aeruginosa* mutants. *Gene* **212**, 77–86 (1998).
13. M. L. Asikyan, J. V. Kus, L. L. Burrows, Novel Proteins That Modulate Type IV Pilus Retraction Dynamics in *Pseudomonas aeruginosa*. *Journal of Bacteriology* **190**, 7022–7034 (2008).
14. D. C. K. Chan, L. L. Burrows, *Pseudomonas aeruginosa* FpvB Is a High-Affinity Transporter for Xenosiderophores Ferrichrome and Ferrioxamine B. *mBio* **14**, e03149-22 (2022).
